# Low lamin A levels enhance confined cell migration and metastatic capacity in breast cancer

**DOI:** 10.1101/2021.07.12.451842

**Authors:** Emily S. Bell, Pragya Shah, Noam Zuela-Sopilniak, Dongsung Kim, Alice-Anais Varlet, Julien L.P. Morival, Alexandra L. McGregor, Philipp Isermann, Patricia M. Davidson, Juniper J. Elacqua, Jonathan N. Lakins, Linda Vahdat, Valerie M. Weaver, Marcus B. Smolka, Paul N. Span, Jan Lammerding

**Author notes:** Correspondence: Jan Lammerding, Department of Biomedical Engineering Cornell University, 235 Weill Hall, Ithaca, NY 14853-7202. The authors declare no competing financial interests.

## Abstract

Aberrations in nuclear size and shape are commonly used to identify cancerous tissue. However, it remains unclear whether the disturbed nuclear structure directly contributes to the cancer pathology or is merely a consequence of other events occurring during tumorigenesis. Here, we show that highly invasive and proliferative breast cancer cells frequently exhibit Akt-driven lower expression of the nuclear envelope proteins lamin A/C, leading to increased nuclear deformability that permits enhanced cell migration through confined environments that mimic interstitial spaces encountered during metastasis. Importantly, increasing lamin A/C expression in highly invasive breast cancer cells reflected gene expression changes characteristic of human breast tumors with higher *LMNA* expression, and specifically affected pathways related to cell-ECM interactions, cell metabolism, and PI3K/Akt signaling. Further supporting an important role of lamins in breast cancer metastasis, analysis of lamin levels in human breast tumors revealed a significant association between lower lamin A levels, Akt signaling, and decreased disease-free survival. These findings suggest that downregulation of lamin A/C in breast cancer cells may influence both cellular physical properties and biochemical signaling to promote metastatic progression.

## Introduction

Abnormalities in the size and shape of cell nuclei have long been recognized as consistent and pervasive features across many types of cancer; accordingly, these features are routinely used in cancer diagnosis and prognosis [1]. However, this predictive power does not distinguish whether nuclear shape abnormalities have direct functional contributions in cancer progression or simply serve as morphological biomarkers. The nucleus is the largest and stiffest organelle, measuring ≈2- to 10-times stiffer than the cell cytoplasm [2–5]. Consequently, deformation of the nucleus can limit cell movement through confined environments [6], including interstitial spaces *in vivo*, where passageways range from 0.1 to 30 μm in diameter [7–9]. Increased nuclear deformability facilitates transit through confined environments [10–13]. Thus, cancer-associated nuclear alterations that result in softer nuclei could favor tumor invasion and metastasis.

Altered nuclear morphology can be driven by changes in chromatin organization [14, 15] or perturbed expression of nuclear envelope (NE) proteins [16–18]. The NE separates the nuclear interior from the cytoplasm and includes the inner and outer nuclear membranes, nuclear pore complexes, and the nuclear lamina, which is a network of lamin intermediate filaments underlying the inner nuclear membrane [19]. In mammalian somatic cells, the nuclear lamina is comprised primarily of A-type lamins (lamins A and C), which are alternatively spliced from the *LMNA* gene, and B-type lamins (lamin B1 and B2), which are encoded by the *LMNB1* and *LMNB2* genes. A-type lamins are the main determinants of nuclear stiffness for large deformations, whereas chromatin structure and composition govern nuclear stiffness during small scale nuclear deformations [20–25]. Accordingly, A-type lamins play an important protective role for stabilizing the nucleus in mechanically-stressed cells, such as in skeletal and cardiac tissues [23, 26–28] and in cancer cells [29], particularly when migrating through confined 3D environments [11, 30, 31] and resisting fluid shear stress in circulation [32].

In addition to providing mechanical stability, the nuclear lamina acts as a scaffold for interactions guiding the localization and assembly of protein complexes such as transcriptional regulators or other NE proteins, tethering and transcriptional silencing of regions of chromatin, and connecting the nuclear interior to the cytoskeleton via the Linker of Nucleoskeleton and Cytoskeleton (LINC) complex [33–35]. A critical role for lamin A/C in tissue function is supported by the multitude of mutations throughout the *LMNA* gene that cause human diseases [36, 37]. Since lamins regulate diverse cellular characteristics, it is striking, but perhaps not surprising, that changes in lamin A/C expression have been documented in many types of human cancer [16, 17, 38], including breast cancer [39–42]. These studies have established links between lamin alterations and clinicopathological features but have not sufficiently addressed the mechanisms of lamin A/C downregulation nor the functional consequences of altered lamin levels on cancer cell biology. Given the prevalence of lamina alterations in tumors and the complex nature of the network of lamin-regulated cellular processes, there is a critical need to establish cause-and-effect relationships for the role of lamins in cancer progression.

In this study, we report that variable expression of lamin A, and to lesser degree lamin C, determines nuclear deformability across a large panel of human and mouse breast cancer lines. Low lamin A levels and increased nuclear deformability were associated with invasive and metastatic breast cancer, increased Akt signaling, and predicted decreased breast cancer survival. Notably, decreased lamin A levels in aggressive breast cancer cells modulated not only nuclear deformability and migration through confined spaces, but also influenced cell morphology, proliferation, and expression of numerous proteins associated with cell-ECM interactions, cell metabolism, and PI3K/Akt signaling. Taken together, our findings suggest that A-type lamins may act as a regulatory node influencing both the biochemical and physical properties of the cell, the dysregulation and downregulation of which promotes breast tumor progression and metastasis through mechanical and non-mechanical functions connected to breast cancer proliferation and migration abilities.

## Materials and methods

### Cells and cell culture

The human breast cancer cell lines MDA-231 (HTB-26), MDA-468 (HTB-132), MCF7 (HTB-22), SKBR3 (HTB-30), HCC70 (CRL-2315), BT-549 (HTB-122), BT-474 (HTB-20), and T47D (HTB-133) were obtained from the American Type Culture Collection (ATCC). MDA-231, MDA-468, MCF7, SKBR3 and T47D cells were cultured in Dulbecco’s Modified Eagle Medium (DMEM) supplemented with 10% (v/v) fetal bovine serum (FBS, Seradigm VWR) and 1% (v/v) penicillin and streptomycin (PenStrep, ThermoFisher Scientific). BT-549, BT-474 and HCC70 cells were grown in Roswell Park Memorial Institute medium (RPMI) 1640 media supplemented with 10% FBS and 1% PenStrep. All cell lines were cultured under humidified conditions at 37°C and 5% CO_2_. The murine epithelial PyMT Flox3 (“PyMT”) tumor cell line was established from an unfloxed tumor arising in the MMTV-PyMT mouse model described in [43] and used as a less metastatic comparator for the highly metastatic Met1 (CVCL_U373) MMTV-PyMT line [44, 45]. PyMT and Met1 cells were maintained in Dulbecco’s Modified Eagle’s Medium: Nutrient Mixture F-12 (DMEM/F12; Gibco) supplemented with 10% (v/v) FBS (Seradigm VWR), and 1% (v/v) penicillin and streptomycin (PenStrep, ThermoFisher Scientific). The 4T1 murine mammary cancer metastatic progression series cell lines (67NR (CVCL_9723), 168FARN (CVCL_0I86), 4T07(CVCL_B383), and 4T1(CVCL_0125))[46] were a kind gift from Dr. Peter Friedl (MD Anderson Cancer Center, Houston, TX, USA). 4T1 progression series cell lines were maintained in RPMI (Gibco) supplemented with L-glutamine (Gibco), sodium pyruvate, 10% (v/v) FBS (Seradigm VWR), and 1% (v/v) penicillin and streptomycin (PenStrep, ThermoFisher Scientific). For PI3K/Akt inhibition experiments, MDA-231 or BT-549 cells were treated with 2 μM NVP-BKM120 (Cayman Chemical, CAS #944396-07-0), 5 μM of Afuresertib (GSK2110183, Selleckchem, Cat #S7521), or vehicle control (DMSO, Sigma, Cat #D2650) for 24 or 48 hours before analysis. For IPTG-induced knockdown of *Lmna* in PyMT cells, IPTG was applied at a concentration of 0.25 mM for at least 48 hours prior to the start of experiments. None of the cell lines in this study are listed NCBI Biosample database of commonly misidentified cell lines. The identities of the BT-549 and MDA-231 cell lines were verified using ATCC STR profiling services. All cell lines were tested for mycoplasma at the beginning of the studies (tested negative with MycoSensor PCR Assay Kit, Agilent, #302109) and have never exhibited contamination symptoms after initial testing.

### Generation of stably modified cell lines

MDA-468 cell lines were stably modified with lentiviral vectors to express the nuclear reporter NLS-GFP (pCDH-CMV-NLS-copGFP-EF1-blastiS [30], available through Addgene (#132772)). MDA-468 cells were also modified with lentiviral vectors to stably express shRNA to LMNA and (MISSION anti-LMNA shRNA, available through Sigma, TRCN0000061835, NM_170707.1-752s1c1) or control non-target MISSION control shRNA (available through Sigma). MDA-231 and BT-549 cells were modified with NLS-RFP (pCDH-CMV-3xNLS-TagRFP-T-EF1-blastiSBT-549 [30]) and then additionally modified with lentiviral vectors to express Lamin A (pCDH-CMV-hLamin_A-IRES-copGFP-EF1-puro [27], available through Addgene (#132773)) or the control plasmid, pCDH-CMV-IRES-copGFP-EF1-puro, generated in house). For IPTG-inducible *Lmna* knockdown, PyMT cells were modified with pLKO.1-EEF1a-mApple-Luc2 and pLKO.1 U63xlacIbs [47] containing shLmna (Sense: GCTTGACTTCCAGAAGAACAT) or shGFP (Sense: GCAAGCTGACCCTGAAGTTCAT) shRNA.

For viral modifications, pseudoviral particles were produced as described [48]. In brief, 293-TN cells (System Biosciences, SBI) at 70-80% confluency were co-transfected with the lentiviral plasmid and lentiviral helper plasmids (psPAX2: Addgene #12260 and pMD2.G: Addgene #12259, gifts from Didier Trono) using PureFection (SBI System Biosciences) following the manufactures protocol. Lentivirus containing supernatants were collected at 48 hours and 72 hours after transfection and filtered through a 0.45 µm syringe filter. Cells were transduced for 1-3 consecutive days with the viral stock diluted 1:3 with culture media in the presence (or absence for BT-549 cell line) of 8 µg/mL polybrene (Sigma-Aldrich). The viral solution was replaced with fresh culture medium, and cells were cultured for 24-48 hours before selection with 1 μg/mL of puromycin (InvivoGen) or 6 µg/mL of blasticidine S (InvivoGen) for 10 days. Cells were also sorted using fluorescence assisted cell sorting (FACS) (BD FACSARIA) to ensure a consistent level of expression of the fluorescent reporters.

### Fabrication and use of microfluidic migration devices

Confined migration experiments in microfluidic devices were performed as described previously [10, 30, 49]. Briefly, the devices contain channels of 5 µm height with constriction widths ranging from 1 to 15 µm in width. Unconfined regions of 250 µm height are found on either side of the migration channels region and cells are loaded into one of these regions [49]. Microfluidic devices were produced from molds that were made using two-layered SU-8 photolithography as described previously [10]. Polydimethylsiloxane (PDMS) replicas of the molds were made using a Sylgard 184 PDMS kit (Dow Corning) using the (1:10) manufacturer’s recipe. After devices were cut to size and holes were punched to allow addition of cells and media, a plasma cleaner (Harrick Plasma) was used to covalently bind microfluidic devices to glass coverslips that had been pretreated with 0.2 M HCl, rinsed, and dried. The assembled devices were placed on a 95°C hot plate for 5 min to improve adhesion. Devices were filled with 70% ethanol for sterilization and then rinsed with sterile deionized water. Devices were then coated with 5 µg/ml fibronectin (Millipore) in PBS (MDA-231 or BT-549) or 50 µg/mL Rat Tail Collagen Type I in 0.02 N acetic acid (MDA-468) overnight at 4°C. After the incubation, devices were rinsed with PBS and filled with cell culture medium before loading ∼30,000 cells through the side port into one of the 250 µm height regions. Cells were allowed to attach in a tissue culture incubator for 6-24 hours, depending on the cell line, prior to the start of imaging. Devices were sealed with a glass coverslip to minimize evaporation and were imaged every 10 min for at least 12 hours in phenol-red free DMEM supplemented with 25 mM HEPES and 10% FBS (for MDA-231 and BT-549 cells) or a 0.1%-12% FBS gradient (for MDA-468 cells). Transit times were calculated through manual analysis. Briefly, transit time

### Fabrication and use of micropipette aspiration devices

Micropipette aspiration of cells into 3-µm-wide, 5-µm-high channels was performed using microfluidic devices fabricated as described previously [50]. Standard lithography techniques were used to produce the mask and wafers for the device at the Cornell NanoScale Science and Technology Facility (CNF). PDMS replicas of the microfluidic device molds were cast using Sylgard 184 (Dow Corning). Devices were cut to size and three entrance port holes were punched and the device was mounted onto a glass slide (pretreated with 0.2M HCl) using a plasma cleaner (Harrick Plasma). Devices were filled with PBS containing 2% bovine serum albumin (BSA), 0.2% FBS solution. Cells were trypsinized and resuspended in 2% BSA, 0.2% FBS PBS solution containing 10 μg/ml Hoechst 33342 (Invitrogen) to visualize nuclei. The cell suspension (∼5 × 10^6^ cells/ml) was perfused into the devices at constant pressure using a MCFS-EZ pressure controller (Fluigent). Pressure at the inlet and outlet ports was set to 1.0 and 0.2 pounds per square inch (relative to atmospheric pressure, *P*_atm_), respectively. The remaining port was outfitted with a hand-held pipette to manually backflush cells from the pockets to allow new cells to enter at the start of each image acquisition sequence. Bright-field and fluorescence images were acquired every 5 seconds for 1-2 minutes. Nuclear protrusion length (ΔL) was analyzed for each cell at each time point using a custom-written MATLAB program [50], made available on request.

### Western blot analysis

Cells were lysed in high salt RIPA buffer (50 mM Tris pH 8.0, 150 mM NaCl (or 750 mM for lamins blots to increase solubility), 1% Nonidet P-40, 0.1% SDS, 0.5% sodium deoxycholate) containing freshly added protease (complete EDTA-Free, Roche) and phosphatase (PhosSTOP, Roche) inhibitors. For lamin blots, lysates were vortexed for 5 minutes at 4°C and passed through an insulin syringe five times to shear DNA. Lysates were cleared by centrifugation at 13 000 rpm for 5 minutes at 4°C. Protein concentration was quantified using Bio-Rad Protein Assay Dye, and 25–30 µg of protein lysate was combined with Laemmli buffer and heated for 3 minutes at 95°C. Proteins were separated using an 8%, 10%, or 4–12% polyacrylamide gel with a standard SDS–PAGE protocol. Protein was transferred to a methanol-activated PVDF membrane at room temperature at a voltage of 16V for 1 hour in a semi-dry transfer apparatus. Membranes were blocked using 3% BSA in Tris-buffered saline containing 0.1% Tween-20 (TBST). Primary antibodies were diluted in the same blocking solution and incubated overnight at 4 °C. After three 5-minute rinses in TBST, infrared (IR)-dye conjugate secondary antibodies IRDye 800CW Donkey Anti-Goat (VWR, #102673-336), IRDye 800CW Donkey Anti-Rabbit (VWR, #102673-334) and IRDye 680RD Donkey Anti-Mouse (VWR, #102673-412) were added at 1/5000 in blocking solution for 1 hour at room temperature. After three more TBST washes, membranes were rinsed with water and imaged on an Odyssey CLx imaging system (LI-COR) and quantification was performed in Image Studio Lite (LI-COR). Primary antibodies used for western blots obtained from Santa Cruz Biotechnology, Inc. were Lamin A/C (E-1) (1/1000, AB_10991536), Lamin A/C (N-18) (1/1000, AB_648152), Actin (1/4000, AB_630835), COL6A1(B-4) (1/500, AB_2783834), Fibronectin (EP5) (1/500, AB_627598), p53 (Pab 240) (1/500, AB_628086), and caveolin-1 (4H312) (1/500, AB_1120056). Antibodies were also obtained from the Developmental Studies Hybridoma Bank (DSHB) created by the NICHD of the NIH and maintained at The University of Iowa, Department of Biology, Iowa City, IA 52242. DSHB antibodies were used at a dilution of 0.5 ug/ml and include Lamin A/C (AB_2618203, MANLAC1(4A7) deposited by Morris, G.E.), Tubulin (AB_528499, E7 deposited by Klymkowsky, M.), Actin (AB_528068, JLA20 deposited by Lin, J.J.-C.), HSPB1 (AB_2617267, CPTC-HSPB1-1 deposited by Clinical Proteomics Technologies for Cancer). Additional antibodies used for western blots were Tubulin (1/3000, Sigma, AB_477579), Lamin B1 (1/1000, Abcam, AB_10107828), Histone H3 (1/5000, Abcam, AB_2885180), Akt (1/1000, Cell Signaling Technologies, #4691T), p-Akt(ser473) (1/500, Cell Signaling Technologies, #4060T) HSPB1 (1/1000, Cell Signaling Technologies, #95357), Histone H3 (1/1000, Cell Signaling Technologies, #3638T) and Lamin B1 (1/1000, Proteintech, #12987-1-AP). Blots validating SILAC results and lamin response to Akt inhibition were conducted as above, but with anti-mouse IgG IgM, HRP-linked (Thermo Fisher Scientific, AB_228321) and anti-rabbit IgG HRP-linked (Cell Signaling, AB_2099233) secondary antibodies. Bands were detected using SuperSignal West Femto Maximum Sensitivity Substrate (Thermo Scientific) and the c-Digit imaging system (LI-COR) and analyzed with ImageStudio software (LI-COR).

### Chiaro indentation and analysis

Cells were seeded on fibronection-coated 35-mm glass bottom dishes (Fluorodish FD35-100) at approximately 40% confluency and cultured overnight in a 37°C incubator with 5% CO_2_ until the start of indentations. Prior to indentation, Hoechst 33342 (1:1000 dilution; Invitrogen #H3570) was added to the media and incubated for 10 min to fluorescently label cell nuclei. The Hoechst containing media was replaced with fresh cell culture media prior to the indentation experiments. Evaluation of nuclear stiffness was performed using a nanoindentation device (Chiaro, Optics11, Netherlands) mounted on an inverted epifluorescence microscope (Observer Z1, Zeiss). Measurements were performed with a 0.024 N/m cantilever with a spherical tip of 9 μm radius. The system was calibrated prior to experimentation according to the manufacturer’s instructions and in the appropriate cell culture media. Before each indentation, the probe was initially positioned approximately 5 µm above the apical cell surface. For each indentation, the probe was lowered towards the cell surface at a speed of 700 nm/s by up to 7 μm, held for 1 second, and then retracted to the initial positions. All indentations were performed at room temperature. To determine the Young’s elastic modulus, the contact point was determined using the Optics11 DataViewer software using identical settings for all cells, and a Hertzian model with a Poisson’s Ratio of 0.5 [51] was fitted to the load vs. indentation curve corresponding to the first 1000 nm of indentation. Any indentation with a fit for the Hertzian model below an R^2^ value of 0.85 was excluded from further analysis.

### Human tumor tissue sections and tissue microarray

The relationship between lamin levels and phospho-Akt staining, receptor status and breast cancer patient survival was investigated using a breast tumor tissue microarray (TMA) described in detail in [52]. The TMA consisted originally of samples from 122 patients, and of these 109 tumor cores were undamaged and analyzable for this study. Breast tumor samples were collected from patients who underwent resection surgery of their primary tumors and had not received systemic adjuvant treatment. Patients were treated between January 1991 and December 1996 and data was available from at least 5 years of follow-up or until a recurrence event. Additionally, levels of ER, PR, HER2, and phosphorylated Akt had been previously assessed by immunohistochemistry [53], enabling the comparison of lamin levels across histological subtypes and in tumors negative or positive for phospho-Akt. Immunofluorescence staining, imaging, and quantification was performed while blinded to breast tumor characteristics or patient outcome. All procedures were conducted according to institutional guidelines with informed consent from patients and with institutional approval.

Human tumor sections used for analysis of relationships between proliferation (Ki67 positivity) and lamin levels were obtained from Dr. Linda Vahdat (Weill Cornell Medicine, IRB# 0408007390A002). The 21 paraffin-embedded breast tumors analyzed included 8 HER2-positive tumors and 5 triple-negative tumors and surgery dates ranged from February 1995 to June 2009.

### Immunofluorescence staining

For staining of cell monolayers, cells were cultured on cover slips for 24-48 hours before fixation with 4% PFA in PBS for 20 min at room temperature. Cells were then rinsed in PBS, quenched in 100 mM glycine/PBS for 5 minutes and permeabilized in 0.3% triton/PBS for 10 minutes. Blocking was in 2% BSA in washing buffer (PBS with 0.2% triton X-100 and 0.05% Tween-20) for 1 hour at room temperature. Cells were stained overnight at 4°C with primary antibodies diluted in blocking solutions. After primary antibodies were removed, cells were rinsed three times (5 minutes each) in washing buffer, and fluorescently labeled secondary antibodies were applied in blocking solution for 40-60 minutes. Where applicable, cells were also stained with Alexa Fluor 647 Phalloidin (1/100, Thermo Fisher Scientific, AB_2620155) during secondary antibody incubation. After three more washes in washing buffer, nuclei were counterstained with 4’,6-diamidino-2-phenylindole (DAPI) (Invitrogen, 0.1% stock diluted 1/500 in PBS for use) for 10 minutes prior to rinsing with water and mounting using Hydromount (#17966 from electron Microscopy Sciences).

For staining of paraffin-embedded human breast tumor tissue sections and tissue microarray, sections were deparaffinized in 3 changes of xylenes followed by 100% ethanol, 95% ethanol and 70% ethanol (all 5 min each). Sections were rehydrated with gently running tap water for 5 minutes. For antigen retrieval, sections were transferred to Tris-EDTA, pH9.0 in a coplin jar and placed in a 55°C water bath. After 30 minutes, water bath temperature was increased to 95°C and incubation continued for an additional 20 minutes after reaching 95°C. Water bath was shut off and allowed to cool slowly for 3-4 hours. Each tissue section was circled with a liquid blocker pen and rinsed 5 minutes with PBS. Each section was covered with blocking buffer of 3% BSA and 5% filtered horse serum in PBS-T (PBS with 0.05% triton X-100 and 0.03% Tween-20) for 1 hour. Primary antibodies diluted in blocking buffer were applied overnight in a humidified chamber at 4°C. After primary antibody treatment and three rinses (10 minutes each) with PBS-T, fluorescently labeled secondary antibodies were applied in Block buffer for 1 hour at room temperature followed by three more 10-minute PBS-T washes. Nuclei were counterstained with 4’,6-diamidino-2-phenylindole (DAPI) (Invitrogen, 0.1% stock diluted 1/500 in PBS for use) for 10 minutes prior to rinsing with water and mounting using Hydromount (#17966 from Electron Microscopy Sciences).

Secondary antibodies and dilutions used included Donkey anti-Rabbit Alexa Fluor 488 (1/300, Thermo Fisher Scientific, AB_2535792), Donkey anti-Mouse Alexa Fluor 568 (1/300, Invitrogen, AB_2534013), Donkey anti-Goat Alexa Fluor 647 (1/150, Thermo Fisher Scientific, AB_141844), and Donkey anti-Goat Alex Fluor 568 (1/300, Life Technologies, AB_142581). The primary antibodies and dilutions used for immunofluorescence on cell monolayers were Lamin A/C (E-1) (1/200, Santa Cruz, AB_10991536), Lamin A/C (N-18) (1/50, Santa Cruz Biotechnologies, AB_648152), Lamin A (H102) (1/100, Santa Cruz Biotechnologies, AB_648148), Lamin B1 (1/100, Abcam, AB_10107828), Lamin B1 (1/200 Proteintech AB_2136290), Lamin B (M-20) (1/100, Santa Cruz Biotechnologies, AB_648158), HSPB1 (1/400, Cell Signaling Technologies, #95357), and HSPB1 (2 ug/ml, AB_2617267, CPTC-HSPB1-1 deposited to the DSHB by Clinical Proteomics Technologies for Cancer). Primary antibodies and dilutions used for immunofluorescence on human breast tumor tissues were Lamin A/C (E-1) (1/500, Santa Cruz, AB_10991536), Lamin A (H102) (1/100, Santa Cruz Biotechnologies, AB_648148), Lamin B (M-20) (1/100, Santa Cruz Biotechnologies, AB_648158), Ki67 (1/200, Novus Biologicals, AB_10001977).

### Fluorescence microscopy and image analysis

Microfluidic migration device experiments and immunofluorescence staining images were collected on inverted Zeiss Observer Z1 microscope equipped with temperature-controlled stage (37°C) and CCD camera (Photometrics CoolSNAP KINO) using 20× air (NA = 0.8), 40× water (NA = 1.2), and 63× oil (NA = 1.4) immersion objectives. Airy units for all images were set between 1 and 2.5. The image acquisition for migration experiments was automated through ZEN (Zeiss) software with imaging intervals for individual sections between 5-10 min. Microfluidic micropipette experiments were collected on a motorized inverted Zeiss Observer Z1 microscope equipped with either charge-coupled device cameras (Photometrics CoolSNAP EZ or Photometrics CoolSNAP KINO) or a sCMOS camera (Hamamatsu Flash 4.0) using a 20× air (NA = 0.8) objective. Cells stained for HSP1B by immunofluorescence were imaged using an Olympus IX83 inverted microscope with Slidebook 6.0 software (Intelligent Imaging Innovations), a 60x oil immersion objective (NA = 1.35) and Hamamatsu Orca R2 monochrome CCD camera.

Analysis of immunofluorescence staining of lamins and Ki67 in cell monolayers and human tumors was performed using a custom-written MATLAB program available on request. DAPI-stained nuclei were used for segmentation of individual nuclei. For tumors, regions containing tumor cells, rather than stroma, were manually selected for inclusion in analysis. For each detected nucleus, mean Ki67 fluorescence intensity was calculated, and the nuclear rim was identified as the top 70% (human tumors) or 20% (cell monolayers) of pixels in the lamin staining. Intensity of both A-type and B-type lamins in the nuclear rim were quantified and lamin B signal was used for normalization of lamin A to facilitate comparisons across tumor sections. For Ki67 staining, nuclei were counted as positive if the signal was 2-fold above background for that image. For association of lamin levels and patient survival, the lamin A : lamin B ratios calculated were used to determine the optimal lamin A : lamin B cutoff to divide patients in having a good or bad prognosis (disease free survival). The analysis produced an optimal cutoff of the lamin A:lamin B ratio of 1.983 to define high and low groupings. Differences in survival were assessed using a Mantel-Cox log rank test.

### Proliferation and cell morphology analysis

Cell lines were plated into 96-well plate at an initial density of 500, 1000, or 1500 cells/well and allowed to attach (3-4 hours) at 37°C. Following this, plates were placed into the IncuCyte (Sartorius) incubator microscope system and imaged with a 20× air objective once every 0.5-1 hour for at least 3 days. Four images per well were taken at each time point. Zoom (Sartorius) software was used to establish an image processing mask and generate confluency percentages over time. Doubling time analysis was performed over data range where cells were in exponential growth phase as determined through IncuCyte imaging. Doubling time was calculated using the formula: Doubling time = ln(two)/((ln(CellsT2/CellsT1) / (Time)), where CellsT1 = initial confluency and CellsT2 = final confluency and Time = duration between initial and final counts. Images collected for proliferation experiments in the IncuCyte were also analyzed for cell morphological differences between BT-549 cells with exogenous expression of Lamin A plasmid (+LamA) or control plasmid. Cell spread area and shape analysis was performed using ImageJ [54]. For filopodia analysis, images were blinded and analyzed for number of cells with protrusions. For cross-sectional nuclear area measurements, cells were analyzed in suspension to minimize effects of adherent cell spreading and cytoskeleton dynamics. Trypsinized cells were flowed into the microfluidics micropipette aspiration device at constant pressure and nuclear cross-sectional area was measured for cells that had entered the 10 × 20 μm^2^ pockets. Measurements were conducted using ImageJ [54]. Cells were included in nuclear area analysis regardless of whether they deformed into constrictions at subsequent timepoints. At least 20 cells were analyzed per cell line in each replicate; 3 independent experiments were performed for each cell line.

### Quantitative RT-PCR

For Akt inhibition experiments, MDA-231 cells were treated with 5 μM of Afuresertib (GSK2110183, Selleckchem, Cat #S7521) or DMSO (Sigma, Cat #D2650) for 24 hours before lysis. For analysis of PyMT and Met1 cells, roughly 3 x 10^6^ cells were lysed in 500 ul of Trizol (Ambion, Cat #15596018) for 10 min at room temperature. Chloroform (VWR Cat #0757) was added at a 1:5 ratio, tubes were vigorously shaken and then spun down at full speed for 5 min. The resulting top clear fraction was transferred to a gDNA Eliminator column and RNA extraction was carried out using a RNeasy Plus Mini Kit (Qiagen, Cat #74136) as per manufacturer’s instructions. The cDNA was generated using iScript select cDNA synthesis kit (Bio RAD, Cat #1708897). The qRT-PCR reactions were done using the LightCycler 480 SYBR green I kit (Roche Cat #04077516001), using the following primers at a concentration of 5 μM: Mouse-GAPDH-Frw 5’-GGAGAGTGTTTCCTCGTCCC-3’, mouse-GAPDH-Rev 5’-ATGAAGGGGTCGTTGATGGC-3’, mouse-LMNA-Frw 5’-TCCACTGGAGAAGAGTGC-3’, mouse-LMNA-Rev 5’-CGCTGCAGTGGGAACCA-3’, human-18S-Frw 5’-GGCCCTGTAATTGGAATGAGTC-3’, human-18S-Rev 5’-CCAAGATCCAACTACGAGCTT-3’, human-GAPDH-Frw 5’-TGCGTCGCCAGCCGAG-3’, human-GAPDH-Rev 5’-AGTTAAAAGCAGCCCTGGTGA-3’, human-LMNA-Frw 5’-GACTCAGTAGCCAAGGAGCG-3’, human-LMNA-Rev 5’-TTGGTATTGCGCGCTTTCAG-3’. Reactions were run on the LightCycler 480/384 machine (Roche serial #1263) and analyzed using LightCycler 480 software (v 1.5.0).

### SILAC proteomic analysis

Quantitative proteomic sample preparation and analysis were performed as described previously [55–57]. Briefly, BT-549 cells were cultured for a minimum of three passages (≈2 weeks) in ‘light’ or ‘heavy’ SILAC media as indicated. SILAC media was composed of RPMI 1640 medium for SILAC (Thermo Fisher Scientific) supplemented with 10% (v/v) dialyzed FBS, 2.0 g/L sodium bicarbonate, 105 mg/L L-leucine, 200 mg/L proline, and 1% (v/v) penicillin and streptomycin (PenStrep, ThermoFisher Scientific). The ‘light’ SILAC media was further supplemented with ‘light’ (normal) lysine (^12^C_6_,^14^N_2_) and arginine (^12^C_6_,^14^N_4_), while ‘heavy’ SILAC media was supplemented with ‘heavy’ lysine (^13^C_6_,^15^N_2_) and arginine (^13^C_6_,^15^N_4_) (100 mg/L for each). Cells were grown to 50-70% confluency at the time of sample collection and were collected by washing cells with PBS, briefly scraping to collect cells in PBS, and centrifugation at 1000 rpm for 3 minutes at 4°C, followed by washing the cell pellet an additional time in PBS. Cells were then lysed in modified RIPA buffer lacking SDS (NaF 50 mM, Tris-Cl pH7.5 50 mM, NaCl 150 mM, NP-40 1%, EDTA 5mM, Sodium deoxycholate 0.25%). After 30 minutes on ice, DNA was sheared, followed by centrifugation for 5 minutes at 4°C and protein concentration of the supernatant was determined by Bradford assay. For each experiment (A and B as indicated), equal amounts of protein from BT-549 control and +LamA cells were combined. The proteins were denatured and reduced (1% SDS, 10 mM DTT) and alkylated with iodoacetamide and then precipitated with three volumes of a solution containing 50% acetone, 49.9% ethanol and 0.1% acetic acid. Proteins were trypsin-digested overnight at 37°C in 50 mM Tris-HCl, pH 8 with 150 mM NaCl and 2 M urea. Peptides were acidified (0.2% of trifluoroacetic acid and formic acid) and then purified using a Sep-Pak C18 Vac purification column (Waters). The eluted peptides were resuspended in 80% acetonitrile/1% formic acid and fractionated using Hydrophilic Interaction Chromatography (HILIC). HILIC fractions were injected into a Q-Exactive Orbitrap and the data was analyzed using Sorcerer as described previously [55, 56].

The analysis identified 91 proteins with at least 2-fold difference in expression upon lamin A overexpression. To determine if these results were representative of changes associated with lamin A expression in human breast tumors, we analyzed publicly available transcriptomic expression data from the TCGA database. RNA-seq count matrices from 1150 women with ductal and lobular neoplasms primary tumors were downloaded from the TCGA database. Count matrices were then pasted together into one matrix and loaded into R (Version 4.1.1) [58]. Using the DESeq2 R package (Version 1.132.0) [59], counts were first subjected to variance-stabilizing transformation, resulting in a log-based transformation and library size normalization of each sample. Linear correlation plots between normalized gene counts were generated using the ggpubr R package (Version 0.4.0) [60] and the Pearson correlation setting. To create *LMNA* “high” and “low” groups, samples with normalized *LMNA* counts one standard deviation above (≥15.32, *n* = 153) or below (≤13.73, *n* = 145) the mean, were assigned into *LMNA* “high” or “low” groups, respectively. DESeq2 was then performed on these two groups to identify differentially expressed genes (DEGs), which were filtered for adjusted *p*-value of *p* < 0.05. Genes were considered “overlapping” if both transcriptomic and proteomic (identified through the SILAC study) differential expression were in the same direction (i.e., both were either up- or down-regulated). To determine if this overlap was statistically significant, the 298 samples previously used in the *LMNA* “high” and “low” groups were instead randomly assigned to groups, and differential gene expression analysis was performed on these randomly assigned groups and filtered based on the same criteria. This random sampling and subsequent differential analysis was performed 1000 times, and the number of identified DEGs “overlapping” genes matching directional regulation in the +Lam A proteomics dataset was analyzed to determine the overall distribution of results.

The 91 proteins identified with at least 2-fold difference in expression between the lamin A overexpressing BT-549 cells and mock controls were further analyzed by STRING network analysis [61]. A protein network was constructed using a confidence interval of 0.6 and proteins with no network connections were removed prior to identification of significantly enriched KEGG pathways against a reference dataset of all proteins identified by mass spectrometry. For upstream regulator identification, proteomic SILAC data was also analyzed with the use of QIAGEN Ingenuity Pathway Analysis (IPA) (QIAGEN Inc., https://digitalinsights.qiagen.com/IPA) [62]. Log-fold change (>2) values were used as inputs for the IPA software. The Upstream Regulator Analysis module was then used to identify potential transcriptional regulators. An activation z-score >1.5 and <–1.5 and Benjamini-Hochberg–corrected *p*-value <0.05 were used to identify significant upstream regulators in our dataset.

### Statistical analysis

Unless otherwise noted, all values are expressed as mean ± standard error of the mean (SEM), and data were generated from a minimum of two independent biological replicates. In figure legends, *N* is used to indicate the number of independent biological replicate experiments, and *n* is used to indicate the number of cells analyzed. Values were considered significantly different for *p* < 0.05. The statistical tests were performed using Prism software (Graphpad) or SPSS (IBM). Normality and variance were assessed in Prism and statistical tests and corrections are indicated in each figure legend. For correlations shown in Figure 1 (F-G, I-L) and Figure S1 (E, G-I), there was no significant departure from linearity based on runs test and similar results could be obtained by either Pearson correlation (linear regression p values shown) or Spearman correlation.

**Figure 1.**
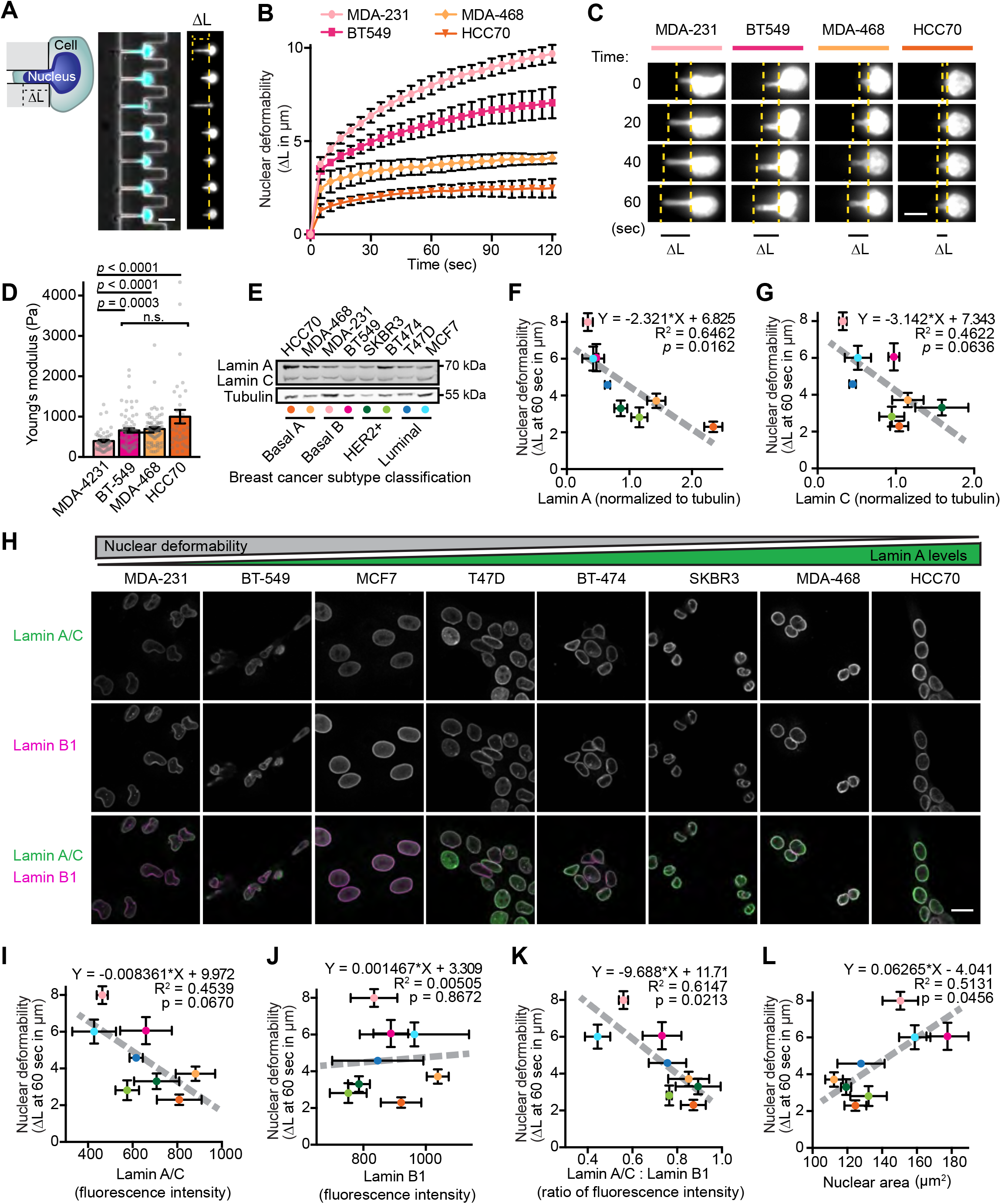
Nuclear size and stiffness vary widely across breast cancer cell lines and correspond to lamin A/C levels. **(A)** Images and schematic of nuclear deformation into a microfluidic micropipette aspiration device under an externally applied pressure. The length of nuclear protrusion (ΔL) into the 3 × 5 μm^2^ channel was measured every 5 seconds at constant pressure application. Scale bar = 20 μm. **(B)** Nuclear protrusion curves for four representative human breast cancer cell lines with different degrees of nuclear deformability. *N* = 3 independent experiments with at least 13 individual measurements per cell line per replicate, mean ± SEM. **(C)** Representative time-lapse image series of deformation of fluorescently labeled nuclei of MDA-231, BT-549, MDA-468, and HCC70 cells in the microfluidic micropipette aspiration device. Scale bar = 10 μm. **(D)** Young’s elastic modulus was determined for the indicated cell lines using a Chiaro nanoindentation device and fitting force-indentations curves to a Hertz model. Data collected across two independent experiments (MDA-231 *n* = 51, BT-549 *n* = 68, MDA-468 *n* = 74, HCC70 *n* = 34). Data displayed as Mean ± SEM and statistical analysis using a Kruskal-Wallis test with Dunn’s multiple comparisons. **(E)** Representative example of Western blot for lamin A/C in a panel of human breast cancer cell lines. Breast cancer subtype classifications are indicated; colored dots correspond to the data for each cell line presented in panels F-G. Tubulin served as loading control. **(F and G)** Nuclear deformability shows an inverse correlation with lamin A (**F**) and lamin C (**G**) levels determined by western blot analysis (*N* = 3). Mean ± SEM. Deformability measurements were taken as nuclear protrusion (ΔL) after 60 seconds of micropipette aspiration in a microfluidic device (*N* = 3 independent experiments, 5-61 measurements per cell type per replicate as shown in Figure S1C). The linear regression results are indicated on each graph. **(H)** Representative examples depicting lamin A/C and lamin B1 immunolabeling for each human breast cell line studied. Scale bar = 20 μm. **(I-K)** Quantification of lamin intensity at the nuclear rim by immunofluorescence (*N* = 3 independent experiments with a minimum of 46 measurements per cell line in each experiment) reveals an inverse correlation between nuclear deformability and A-type lamin levels or lamin A/C: lamin B1 ratio, but not for lamin B1 alone. Deformability measurements same as in panels F-G. Data are plotted as mean ± SEM; the linear regression results are indicated on each graph. **(L)** Larger nuclear cross-sectional area correlates with increased nuclear deformability. Quantification of nuclear cross-sectional area of trypsinized cells in suspension (*N* = 3 independent experiments with a minimum of 20 measurements per cell line per replicate, Mean ± SEM). Deformability measurements same as in panels F-G. Linear regression analysis is indicated on each graph.

## Results

### Invasive breast cancer cells exhibit increased nuclear deformability correlating with lower A-type lamin levels

Breast cancer is a heterogeneous disease that can be categorized into subtypes with distinct prognoses and treatment options [63, 64]. To investigate the role of nuclear deformability in breast cancer pathogenesis, we used a novel microfluidic micropipette aspiration assay [50] to assess nuclear mechanics in a panel of breast cancer cell lines reflecting the diverse molecular and cellular phenotypes of breast cancer (Fig. 1 A, Fig. S1 A, Video 1, and Table S1). Nuclear deformability was heterogeneous within cell lines and varied dramatically across different cell lines (Figs. 1 B and C, Fig. S1 B and C), suggesting that some lines harbor intrinsic changes that influence nuclear mechanical properties. To provide an independent measure of nuclear deformability, we conducted microindentation experiments on a subset of cell lines. These experiments revealed a similar trend in nuclear deformability, with MDA-MB-231 cells exhibiting the softest nuclei, and HCC70 cells the stiffest nuclei (Fig. 1D and Fig. S1D). The differences in nuclear deformability were more pronounced in the micropipette aspiration experiments, likely due to the substantially larger nuclear deformations achieved (2-8 μm deformation vs. 1 μm indentation), which may also reflect that nuclear mechanical responses are controlled by different factors (i.e. nuclear lamina vs. chromatin) at large scale vs. small scale deformations [20]. Thus, for subsequent measurements of nuclear deformability, we used the micropipette aspiration assay. Importantly, the three cell lines (MDA-MB-231, hereafter MDA-231, BT-549, and MCF7) with the most deformable nuclei have all been previously characterized to be invasive *in vitro*, fast growing, and amenable to use in metastasis assays in immunocompromised mice (Table S1), suggesting that increased nuclear deformability could be a characteristic feature of metastatic breast cancer cells. Since lamin A/C are major determinants of nuclear deformability [24], we assessed lamin levels in the panel of cell lines by Western blot analysis and immunofluorescence. Since lamins A and C share the first 566 aa and vary only in their C-terminus, most antibodies recognize both lamin A and C; thus, for this immunofluorescence analysis, we collectively quantified lamin A/C levels. Cell lines with highly deformable nuclei had decreased lamin A and C levels, and this association was stronger for lamin A than for lamin C (Figs. 1 E-I). Interestingly, decreased ratios of lamin A to lamin C also associated with increased nuclear deformability, though this association did not reach statistical significance (Fig. S1 E). Unlike lamin A/C, decreased lamin B1 levels did not correlate with increased nuclear deformability (Fig. 1 I-J), but the ratio of lamin A/C to lamin B1 by immunofluorescence showed a strong inverse correlation with nuclear deformability (Fig. 1 K). This finding supports the use of lamin B1 as a reliable normalization signal when assessing A-type lamins by immunofluorescence, in accordance with the relatively consistent levels of B-type lamins observed across tissues, developmental stages, and tumorigenesis [23, 65, 66], although we cannot exclude the possibility that the lamin A/C : lamin B1 ratio may directly govern nuclear deformability [23]. Cells with increased deformability and decreased lamin A/C levels or lamin A/C:B1 ratios also exhibited larger cross-sectional nuclear area (Figs. 1 L, Fig. S1 F-H), whereas lamin B1 levels alone showed no significant correlation with nuclear area (Fig. S1 I). As enlarged nuclei are a feature of advanced grade cancers and worse prognosis [67], these data further support an association between decreased lamin A/C levels and advanced, aggressive breast cancer.

### A-type lamin expression is a major determinant of breast cancer nuclear deformability

In addition to the composition of the nuclear lamina itself, nuclear deformability is influenced by chromatin modifications [20, 21, 68] and contributions from the actin cytoskeleton [69, 70]. Since the nuclear lamina is connected to both heterochromatin formation and cytoskeletal organization [33–35], alterations in lamin A could influence nuclear deformability through both mechanical properties of the lamina itself and altered regulation of chromatin and cytoskeleton. To determine how each of these factors may contribute to the nuclear deformability characteristics observed across breast cancer cell lines, we used the MDA-468 cell line, which had low nuclear deformability and high lamin A/C levels (Fig. 1 B, F-H, Fig. S1 B). Nuclear deformability was assessed by micropipette aspiration following shRNA-mediated depletion of lamin A/C, decreasing heterochromatin through inhibition of histone deacetylases with Trichostatin A (TSA), and disruption of actin polymerization using Cytochalasin D (CytoD), and compared to non-target and vehicle controls (Fig. 2 A and B). As expected, depletion of lamin A/C resulted in a large (2.3-fold) increase in nuclear deformability compared to non-target shRNA controls (Fig. 2 C). In contrast, disrupting actin filaments with CytoD resulted in only a mild (1.4-fold) increase in nuclear deformability, and increasing euchromatin levels with TSA did not significantly change nuclear deformability (Fig. 2 C), indicating that direct alterations in nuclear lamina composition and mechanics are the major determinants of the variable nuclear stiffness observed with differential lamin A/C expression across the panel of breast cancer cells. In further support of this idea, increasing lamin A levels by ≈1.5-2.5-fold (Fig. 2 D, E, G, and H) through stable exogenous expression in two highly invasive breast cancer cell lines, BT-549 and MDA-231, which originally had low lamin A/C levels and very deformable nuclei, resulted in significantly reduced nuclear deformability (Fig. 2 F and I).

**Figure 2.**
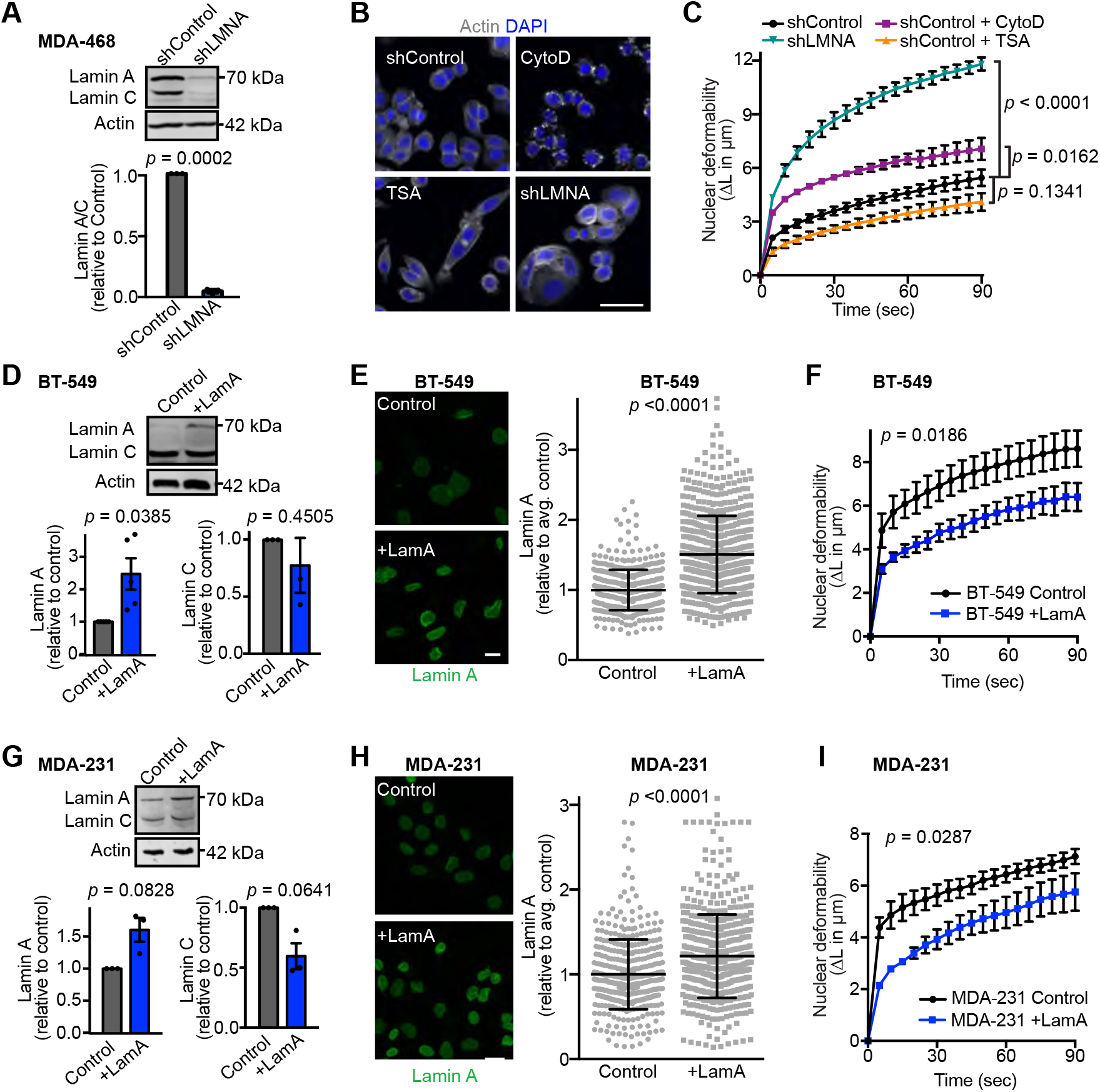
Decreased A-type lamin levels result in the enhanced nuclear deformability of invasive breast cancer cells. **(A)** Representative western blot and quantification (*N* = 3, mean ± SEM) showing decreased lamin A/C levels in MDA-468 cells expressing *LMNA* shRNA or non-target control. Lamin A/C levels were normalized to non-target control in each Western blot. Significance based on one-sample *t* test compared to a theoretical value of 1. (**B**) Representative images of MDA-468 cells stained for DNA (DAPI) and F-actin (phalloidin) to assess the effect of lamin A/C depletion (shLMNA), inhibition of actin polymerization by cytochalasin D (CytoD, 4 μM for 20 minutes), or inhibition of histone deacetylation by trichostatin A (TSA, 4 nM for 12 hours) on cell and nuclear morphology. Scale bar = 20 μm. (**C**) Micropipette aspiration in a microfluidic device was used to assess which cellular components have the greatest impact on nuclear deformability. MDA-468 shLMNA cells and MDA-468 and non-target controls (shControl) cells treated with CytoD, TSA, or DMSO vehicle control. Statistical analysis based on two-way repeated measures (RM) ANOVA with Tukey’s multiple comparisons test. *N* = 3 independent experiments with at least 34 individual measurement per condition, per replicate, mean ± SEM. (**D**) Representative Western blot and corresponding quantification from five independent experiments probed for lamin A/C lamin levels in BT-549 cells with exogenous expression of Lamin A (+LamA) and mock controls. Actin was used as loading control. Statistical analysis based on one-sample *t* test with a theoretical value of 1 (control). Data shown as mean ± SEM. (**E**) Examples of immunofluorescence staining of lamins in BT-549 cells expressing exogenous lamin A (+LamA) or control construct and quantification of nuclear rim lamin A staining intensity. Statistics based on two-tailed Mann-Whitney test. Mean ± SD, *n* = 475 and 661 cells quantified across 3 independent experiments. Scale bar = 20 μm. (**F**) BT-549 cells with addition of lamin A exhibit decreased nuclear deformability as determined by micropipette aspiration in a microfluidic device. Statistical analysis based on two-way RM ANOVA. Data depicted as mean ± SEM for *N* = 7 independent experiments with at least 6 individual measurement per cell per replicate, for a total of 153 control and 170 +LamA measurements. (**G**) Representative western blot for lamin A in MDA-231 control and +LamA cells and corresponding quantification based on three independent experiments. Statistical analysis based on one-sample *t* test with a theoretical value of 1. Data plotted as mean ± SEM. (**H**) Examples of immunofluorescence staining of lamins in MDA-231 cells expressing exogenous Lamin A (+LamA) or control construct and quantification of nuclear rim lamin staining by immunofluorescence. Statistical analysis based on two-tailed Mann-Whitney test. Mean ± SD, *n* = 404 and 546 cells quantified across 2 independent experiments. Scale bar = 20 μm. (**I**) MDA-231 cells with addition of lamin A exhibit decreased nuclear deformability as determined by micropipette aspiration in a microfluidic device. Statistical analysis based on two-way RM ANOVA. Data depicted as mean ± SEM for *N* = 3 independent experiments with at least 5 individual measurement per cell per replicate, for a total of 120 Control and 103 +LamA measurements.

### Decreased Lamin A levels and increased nuclear deformability facilitate migration through confined spaces

During progression through the metastatic cascade, cells disseminating from the primary site encounter various passageways substantially smaller than the diameter of the nucleus, such as during invasion through the basement membrane, migration through tight interstitial spaces, or during intra- and extravasation [7–9]. Deformation of the nucleus to squeeze through such confined spaces can impose a rate-limiting step on cell passage [6, 71]. Therefore, loss of A-type lamin levels may contribute to metastatic progression by increasing nuclear deformability and enhancing migration through confined environments. To test this hypothesis in isogenic cell line models, we assessed whether depletion of lamin A/C by shRNA enhanced confined migration in cells with high lamin A/C levels, and conversely whether exogenous expression of lamin A in cells with low lamin A/C levels impaired their migration through tight spaces. To precisely control the degree of confinement, we used microfluidic migration devices that contain small (1×5 μm^2^ and 2×5 μm^2^) constrictions, as well as larger (15×5 μm^2^) control channels that do not require nuclear deformation (Fig. 3A)[10, 30, 49]. MDA-468 cells depleted for lamin A/C migrated significantly faster through small constrictions than non-target control cells; in contrast, lamin A/C depleted and non-target controls had similar transit times in the larger control channels (Fig. 3 B and C, Video 2), indicating that loss of lamin A/C specifically enhanced migration through confined spaces. Conversely, exogenous expression of lamin A in BT-549 and MDA-231 cells, which normally have low lamin A/C expression and high nuclear deformability (Fig. 1B-G), resulted in decreased nuclear deformability (Figs. 2 F and I) and slower transit through small constrictions relative to control cells (Fig. 3 D-G, Video 3), indicating that nuclear deformability acts as a critical determinant of confined migration efficiency even in highly invasive cancer cells. In contrast, the effect of lamin A overexpression on cell transit times was much less pronounced in the larger control channels, which overall allowed faster cell transit (Figs. 3 E and G). Taken together, these results highlight that breast cancer cells with diminished lamin A/C levels and corresponding highly deformable nuclei have a migration advantage specifically in confined environments, which may facilitate invasion and metastatic progression.

**Figure 3.**
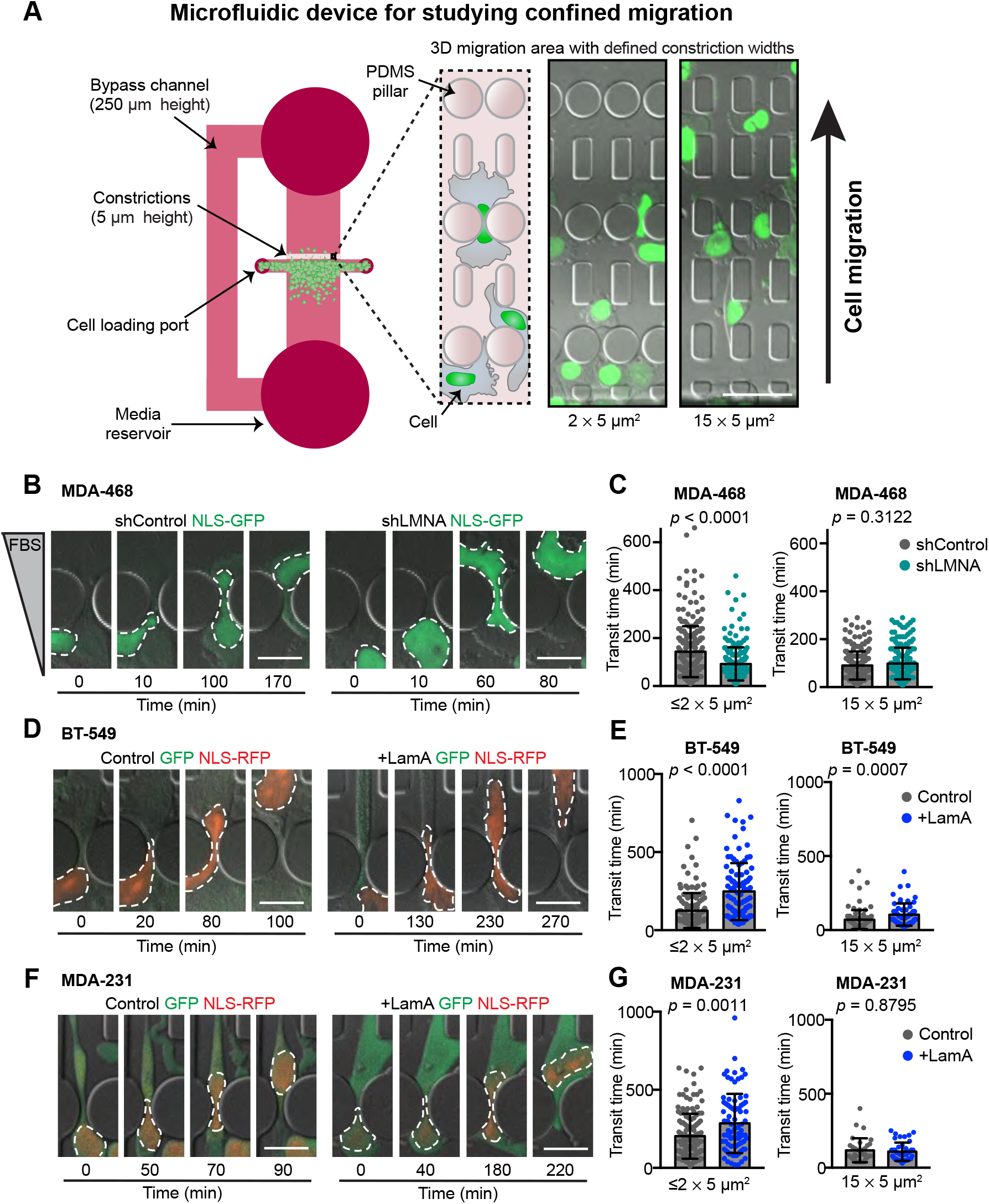
Decreased Lamin A levels facilitate migration through confined spaces. (**A**) Schematic of the PDMS microfluidics devices used for analysis of cell migration through confined 3-D environments. Cells are plated into the 250 μm tall space in front of a 5 μm tall migration area containing channels with constrictions that are either 1 × 5 μm^2^, 2 × 5 μm^2^, or 15 × 5 μm^2^ in size. For MDA-468 cells, a chemoattractant of 0.1% to 12% FBS gradient was used to stimulate migration experiments, but a gradient was not needed to stimulate movement of MDA-231 or BT-549 cells. Cells are imaged at 10-minute intervals. Transit times are quantified based on the movement of the fluorescently labeled nucleus through the constrictions. Representative images show NLS-GFP-expressing MDA-468 cells ≈20 hours post-seeding in the device. Scale bar = 50 μm. **(B)** Representative image sequences of MDA-468 cells expressing shControl and shLMNA as the cells migrate through ≤2 × 5 μm^2^ constrictions along a 0.1% to 12% FBS gradient. (**C**) Transit times for MDA-468 shControl and shLMNA cells moving through ≤2 × 5 μm^2^ (*n* = 252 and 213 cells) or 15 × 5 μm^2^ (*n* = 224 and 206 cells) constrictions along a 0.1% to 12% FBS gradient. (**D**) Representative image sequences of BT-549 Control and +LamA cells migrating through ≤2 × 5 μm^2^ constrictions. (**E**) Transit times for BT-549 Control and +LamA cells moving through ≤2 × 5 μm^2^ (*n* = 142 and 108 cells) or 15 × 5 μm^2^ (*n* = 71 and 61 cells) constrictions. (**F**) Image sequences of MDA-231 Control and +LamA cells migrating through ≤2 × 5 μm^2^ constrictions. (**G**) Transit times for MDA-231 Control and +LamA cells moving through ≤2 × 5 μm^2^ (*n* = 142 and 83 cells) or 15 × 5 μm^2^ (*n* = 33 and 42 cells) constrictions. For all cell lines, transit time calculations were collected from across a minimum of three independent experiments and displayed as mean ± SD. Statistical analysis based on two-tailed Mann-Whitney test. Scale bars = 20 μm.

### Lower lamin A levels are associated with metastatic progression

Our findings support that downregulation of A-type lamins could be a feature of aggressive, metastatic breast cancer. However, a relationship with metastatic progression is difficult to assess solely through comparison of cancer cell lines collected from individuals with different genetic backgrounds and containing distinct molecular mechanisms driving tumor progression. To better assess whether a relationship exists between metastatic progression and downregulation of lamin A/C, we compared lamin levels in isogenic cell lines derived from murine mammary cancer driven by expression of the polyomavirus middle T antigen (PyMT) [72]. The PyMT mammary epithelial cell line was compared with the highly metastatic Met1 variant derived from the same model in a syngeneic background [44]. The metastatic Met1 cells had significantly lower levels of lamin A and lamin C than the primary PyMT line, based on the analysis of protein (Fig. 4 A) and transcript (Fig. 4 B) levels. Accordingly, Met1 cells had significantly more deformable nuclei than the primary tumor PyMT line (Fig. 4 C), and we previously found that Met1 cells exhibited more efficient confined migration in microfluidic device than the primary PyMT line [45]. Depletion of lamin A/C via IPTG-inducible shRNA was sufficient to increase nuclear deformability in the primary PyMT line to approximately that of Met1 cells (Figs. 4 D-F). These findings provide further support that metastatic breast cancers are characterized by increased nuclear deformability driven by decreased A-type lamin levels.

**Figure 4.**
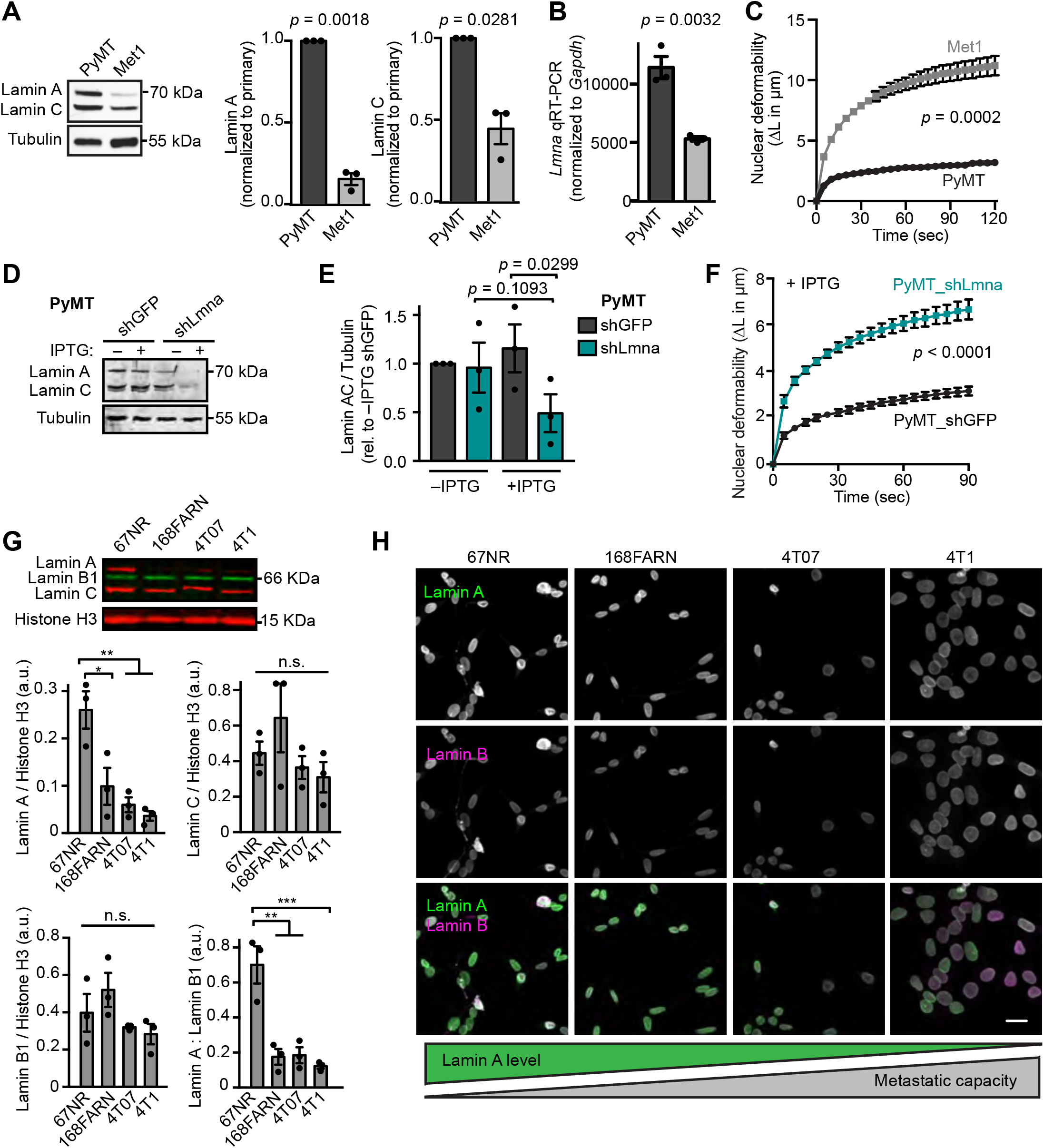
Breast cancer cells with increased metastatic capacity have reduced lamin A expression. **(A)** Representative western blot and corresponding quantification (*N* = 3, mean ± SEM) for lamin A/C levels in MMTV-PyMT transgenic mouse tumor model cell lines from a primary tumor (PyMT) or the highly metastatic Met1 line derived from a late-stage MMTV-PyMT tumor. Tubulin served as a loading control. Statistical analysis based on one-sample *t* test with a theoretical value of 1. **(B)** Quantitative RT-PCR for *Lmna* mRNA in the metastatic Met1 cells compared to the primary tumor PyMT line. Mean ± SEM, *N* = 3. Statistical analysis by two-tailed unpaired Student’s *t* test. **(C)** Nuclear protrusion curves comparing nuclear deformability of Met1 and PyMT cells upon micropipette aspiration in a custom microfluidic device. Mean ± SEM, *N* = 3, independent experiments with at least 10 individual measurements per cell line, per replicate. Statistical analysis based on two-way RM ANOVA. **(D-E)** Western blot and corresponding quantification (*N* = 3, mean ± SEM) of lamin A/C levels in PyMT cells expressing IPTG-inducible shRNA to *Lmna* or non-targeting (shGFP) control. Cells were cultured in media with or without 0.25 mM IPTG for 48 h prior to analysis. Statistical analysis based on one-way ANOVA with Tukey’s multiple comparisons test. Tubulin was used as a loading control. **(F)** PyMT cells depleted for A-type lamins as in (D) and (E) exhibit increased nuclear deformability as quantified by micropipette aspiration in a microfluidic device. *N* = 4 independent experiments with at least 9 individual measurements per cell type, per replicate for a total of 111 shGFP and 142 sh*Lmna* measurements. Data depicted as mean ± SEM. Statistical analysis based on two-way RM ANOVA. **(G)** Representative western blot and corresponding quantification (*N* = 3, mean ± SEM) of lamin levels in cell lines from the 4T1 mouse mammary tumor metastatic progression series. Histone H3 is included as a loading control. Statistical analysis based on one-way ANOVA with Tukey’s multiple comparisons. *, *p* = 0.0192, **, *p* <0.01, ***, *p* = 0.0008. **(H)** Representative examples of immunofluorescence staining of lamins in the 4T1 metastatic progression series cell lines. Cell lines are displayed in order of increasing propensity for metastasis during tumor growth in the mouse mammary fat pad. Schematic representation of lamin levels based on the general trend shown in panel D. Scale bar = 20 μm.

The above findings suggest that decreased expression of A-type lamins may provide a selective advantage at some point during breast cancer metastasis. Metastasis is a multi-step process involving local invasion, intravasation, survival in circulation, arrest in vasculature, extravasation, and secondary tumor growth [73]. To evaluate which stage of metastasis is associated with the decrease in A-type lamins, we analyzed lamin expression in the isogenic 4T1 progression series. These sister cell lines were all isolated from a single, spontaneously arising tumor from a Balb/cfC3H mouse and are each capable of progressing to different stages of metastasis [46, 74]. Strikingly, increased metastatic capacity was associated with a significant decrease in lamin A levels. The 67NR cell line, which does not leave the primary site, had the highest lamin A levels, whereas cells capable of disseminating to the lymph nodes (168FARN cells) and lungs (4T07) had significantly lower lamin A expression, and lamin A levels were lowest in cells capable of metastasizing to the lungs and bone (4T1) [46, 75] (Fig. 4 G and H). In contrast, lamin B1 and lamin C levels did not vary significantly across the progression series (Fig. 4 G). These findings support that a decrease in lamin A levels is acquired or selected for during the early invasion or intravasation stages of the metastatic cascade.

### Increasing lamin A in breast cancer cells alters mediators of cell-matrix interactions, cell metabolism, and PI3K/Akt signaling

In addition to influencing nuclear deformability and stability, the nuclear lamina participates in diverse cellular processes, including chromatin organization, DNA damage repair, and the regulation of transcription factors and signaling pathways [16, 76]. Consequently, the decrease of lamin A/C in invasive breast cancer may promote tumor progression through mechanisms beyond the altered mechanical properties of the nucleus. To assess additional effects of altered lamin levels on tumor progression in an isogenic model, we performed quantitative proteomic analysis in BT-549 cells stably modified to exogenously express lamin A and mock controls, using stable isotope labeling with amino acids in culture (SILAC) and mass spectrometry analysis (Fig. 5 A). Upon a moderate (≈2.5-fold) increase in lamin A (Fig. 2 D and E), resulting in BT-549 cells with lamin A/C levels similar to breast cancer cell lines with high lamin A/C levels, we identified 91 proteins with at least 2-fold difference in expression between the lamin A overexpressing BT-549 cells and mock controls (Fig. 5 B). Of these, 42 proteins increased in expression with the increase in lamin A levels, and 49 proteins decreased when lamin A was overexpressed. Changes in expression of five of the identified proteins were validated by Western blot analysis (Supp. Fig. S2 A-E). To determine if results from the proteomic analysis in an isogenic cell line model were representative of lamin A-associated gene expression changes in human breast cancer tumors, we analyzed existing transcriptomic dataset of 1150 breast cancer tumors from the TCGA database, and identified 298 tumors as either “*LMNA* high” or *LMNA* low”, based on their *LMNA* expression levels (Supp. Fig. S2 F). Differential gene expression (DEG) analysis between “*LMNA* high” and “*LMNA* low” revealed that of the 91 genes corresponding to the differentially expressed proteins in the SILAC analysis, 48 (53.3%) agreed with the direction of gene regulation (i.e., both were either up- or down-regulated in the SILAC data and tumor RNA-seq data), and 34 of the 48 (37.8% of the total) reached statistical significance in their expression between ‘*LMNA* high’ and ‘*LMNA* low’ groups (Fig. 5 B, Table S2). Notably, a bootstrap analysis using randomly assigned groups confirmed that this overlap was highly significant (*p* < 0.001), as random groupings followed by DEG analysis only yielded a maximum of 11 genes overlapping with the SILAC data in 1000 simulations (Supp. Fig. S2 G). Thus, many of the changes identified upon overexpression of lamin A in the BT-549 breast cancer cell line are also found in human tumors with higher expression of *LMNA*, supporting the idea that lamin A levels may play an integral component in many aspects of tumor biology.

**Figure 5.**
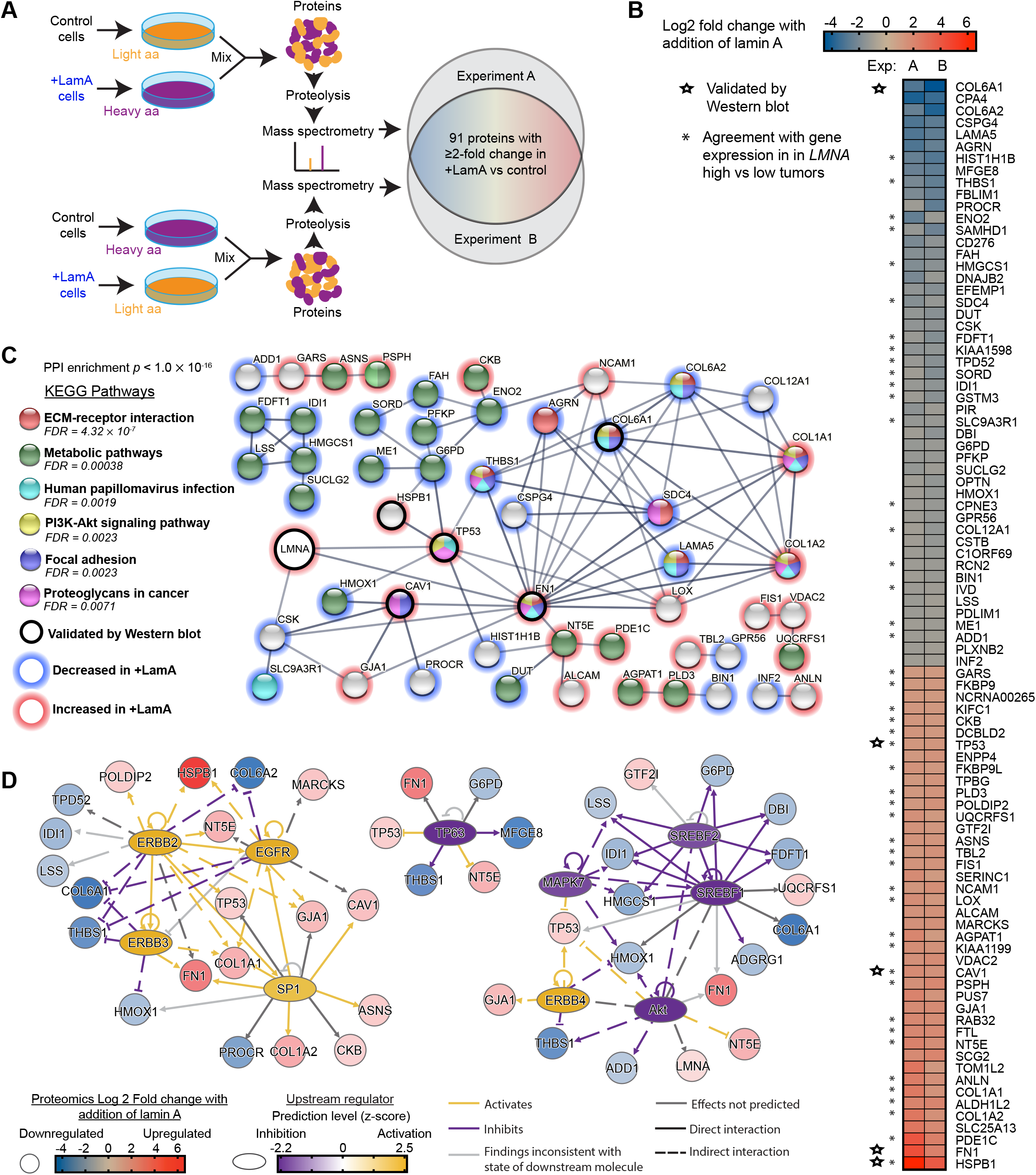
Increasing lamin A in breast cancer cells alters expression of proteins involved in cell metabolism, extracellular remodeling, adhesion, and cytoskeleton dynamics. **(A)** Schematic for the design of the SILAC experiments to identify proteins altered upon increased lamin A expression in BT-549 cells. Cells were cultured in media containing normal/light amino acids or heavy amino acids (L-lysine ^13^C_6_, ^15^N_2_ and L-arginine ^13^C_6_, ^15^N_4_) for 2 weeks prior to analysis. **(B)** Proteins listed here were detected to have ≥ 2-fold change in both of two experiments comparing +LamA and control BT-549 cells. Increase (red) and decrease (blue) in protein abundance in +LamA cells relative to controls is indicated by color on a log2 scale. Combined levels of lamin A and C were detected at ≈1.7-fold increase in +LamA cells compared to control cells in both experiments. Stars indicate proteins selected for validation by western blot analysis. Asterisks indicate proteins that exhibit agreement with direction of differential gene expression in “*LMNA* high*”* compared to low “*LMNA* low” breast tumors. High and low groupings were determined as one standard deviation above (≥15.32, n=153) or below (≤13.73, n=145) the mean for normalized *LMNA* counts, respectively. DESeq2 was performed on these two groups to identify differentially expressed genes (DEGs), which were filtered for adjusted *p*-value < 0.05. **(C)** STRING protein association network analysis (confidence threshold >0.6) for the proteins listed in panel B and *LMNA*. Proteins without any network associations were removed before analysis of KEGG pathway enrichment. Red outlines indicate proteins with increased levels upon lamin A overexpression and blue outlines indicate decreased expression. An additional black outline indicates proteins validated by Western blot. Image adapted from https://string-db.org/. **(D)** Potential upstream transcriptional regulators were identified in the +LamA SILAC proteomic dataset using QIAGEN Ingenuity Pathway Analysis (IPA) (QIAGEN Inc., https://digitalinsights.qiagen.com/IPA). The Upstream Regulator Analysis module was used to identify potential transcriptional regulators. An activation z-score >1.5 and <-1.5 and Benjamini-Hochberg corrected *p*-value <0.05 were used to identify significant upstream regulators. Blue to red coloration indicates decreased or increased protein levels in +LamA relative to control BT-549 cells, and purple to yellow coloration indicates predicted inhibition or activation of potential upstream regulators.

To better understand the cellular pathways affected by lamin expression, *LMNA* and the 91 genes identified by SILAC were assessed using STRING network analysis [61]. A protein-protein interaction (PPI) network (enrichment *p* value of < 0.001) indicated a biological connection between some of the identified proteins (Fig. 5 C). Analysis of the interaction network showed enrichment of KEGG pathways related to interactions between cells and their local environment (“ECM-receptor interactions”, “Focal adhesion”, “Proteoglycans in cancer”), “Metabolite pathways”, and “PI3K-Akt signaling” (Fig. 5 C, Supp. Fig. S2 H). All of these pathways were found to be enriched in the largest interaction network, made up of 35 nodes (38% of total input proteins), including both *LMNA* and highly DEGs like *HSPB1*, *FN1*, and *COL6A1*. While this network appears to be made up of several influential proteins driving these pathway associations, no clear pathway seemed to be dominated by only up- or down-regulated proteins. To further elucidate how these lamin A-associated changes may be related to a potential regulator, we analyzed the list of proteins with at least 2 fold change in the +LamA SILAC dataset by QIAGEN Ingenuity Pathway Analysis (IPA) and the Upstream Regulator Analysis module. This analysis identified SREBF1/2, Sp1, Akt, MAPK7, TP63, EGRBB2/3/4, and EGFR as potential upstream transcriptional regulators, grouped into three mechanistic networks (Fig. 5D, Table S3). These regulators explained most of the lamin A-associated changes observed in the large network identified with STRING, and therefore may play an important role in regulating cellular processes associated with differential Lamin A expression in cancer. Notably, these results include regulators with previously known connections to *LMNA* (i.e., Akt, SREBP1, and SP1) [77–79], further supporting the interpretation that the SILAC approach identified lamin A-associated changes relevant to regulatory interactions in human tumors.

### Increased lamin A expression results in increased HSPB1 levels in breast cancer cell lines and tumors

To further examine the connections between lamin A levels and expression of other proteins in more detail, we examined the protein with the largest increase upon lamin A addition, the molecular chaperone heat shock protein 27 (HSPB1 or HSP27) (Fig. 5 B, bottom). HSPB1 has been implicated in diverse cellular functions, including apoptosis, cytoskeletal dynamics, proliferation, and immune response [80, 81]. Importantly, HSPB1 is normally most highly expressed in cardiac and skeletal muscle [82], which coincides with the critical role of A-type lamins in tissues that experience high levels of mechanical stress and the high expression of A-type lamins in these tissues [23, 83]. HSPB1 was nearly undetectable in control BT-549 cells but increased ≈12-fold upon overexpression of lamin A (Supp. Fig. S2 A). To establish whether A-type lamin expression is a regulator of HSPB1 across breast cancer models, we examined HSPB1 expression in PyMT cells in response to inducible depletion of lamin A/C. Depletion of lamin A/C resulted in significantly decreased HSPB1 levels in the PyMT cells, further supporting our findings in the human breast cancer cells (Supp. Fig S2 I). The relationship between lamin A and HSPB1 also extended to other breast cancer cell lines, as assessment of lamin and HSPB1 levels by western blot revealed a significant positive correlation between HSPB1 and lamin A, but not lamin C (Supp. Fig. S2 J-L), though this association was largely driven by the very high lamin A and HSPB1 levels in HCC70 cells (Supp. Fig. S2 K-L). However, based on the TCGA dataset, HSPB1 was also significantly more highly expressed in “*LMNA* high” breast tumors relative to “*LMNA* low” tumors (Supp. Fig. S2 M) and showed a linear correlation with *LMNA* levels across human breast tumors (Supp. Fig S2 N). Taken together, these data indicate that HSPB1 levels scale with lamin A expression across cell lines, tumors, and species, and support the interpretation that the SILAC proteomics studies identified robust and functionally relevant lamin A-dependent cellular changes in breast cancer cells.

### Increased expression of lamin A and HSPB1 alter tumor cell morphology

Since many of the proteins altered upon changes in lamin A expression are involved in cell matrix remodeling, cell adhesion, and actin dynamics (Table S2), we examined how changes in lamin A/C levels affect cell morphology. BT-549 breast cancer cells in culture are normally characterized by extensive cell spreading and expansive lamellipodia for cell motility (Fig. 6 A). Increasing the expression of lamin A in BT-549 cells resulted in decreased cell spreading and spindle-like cells with some filopodia-like protrusions, corresponding to a dramatic loss of lamellipodia (Figs. 6 A-F), which may contribute to decreased migration abilities (Fig. 3 E). Thus, changes in lamin A expression can impact cellular structure and functions beyond the nuclear envelope, and alteration of lamin A/C expression can lead to numerous proteomic and functional changes within breast cancer cells. Collectively, these changes affect how the cells modify and interact with their physical microenvironment and could promote cancer cell invasion and metastatic progression. To explore if some of these changes could be attributed at least in part to altered expression of HSPB1, we assessed cell morphology and HSPB1 levels in MDA-468 cells by immunofluorescence (Supp. Fig S2 O). Since HSPB1 levels were highly variable within the cell population, we were able to relate levels of HSPB1 with cell morphology across individual cells. Notably, increased HSPB1 expression correlated with decreased spread cell area and increased cell circularity (Supp. Fig. S2 P-Q), suggesting that increased HSPB1 levels could lead to morphologically distinct subset of cells, although other pathways driven by altered lamin A expression (see Fig. 5 C-D) likely contribute to the morphological changes.

**Figure 6.**
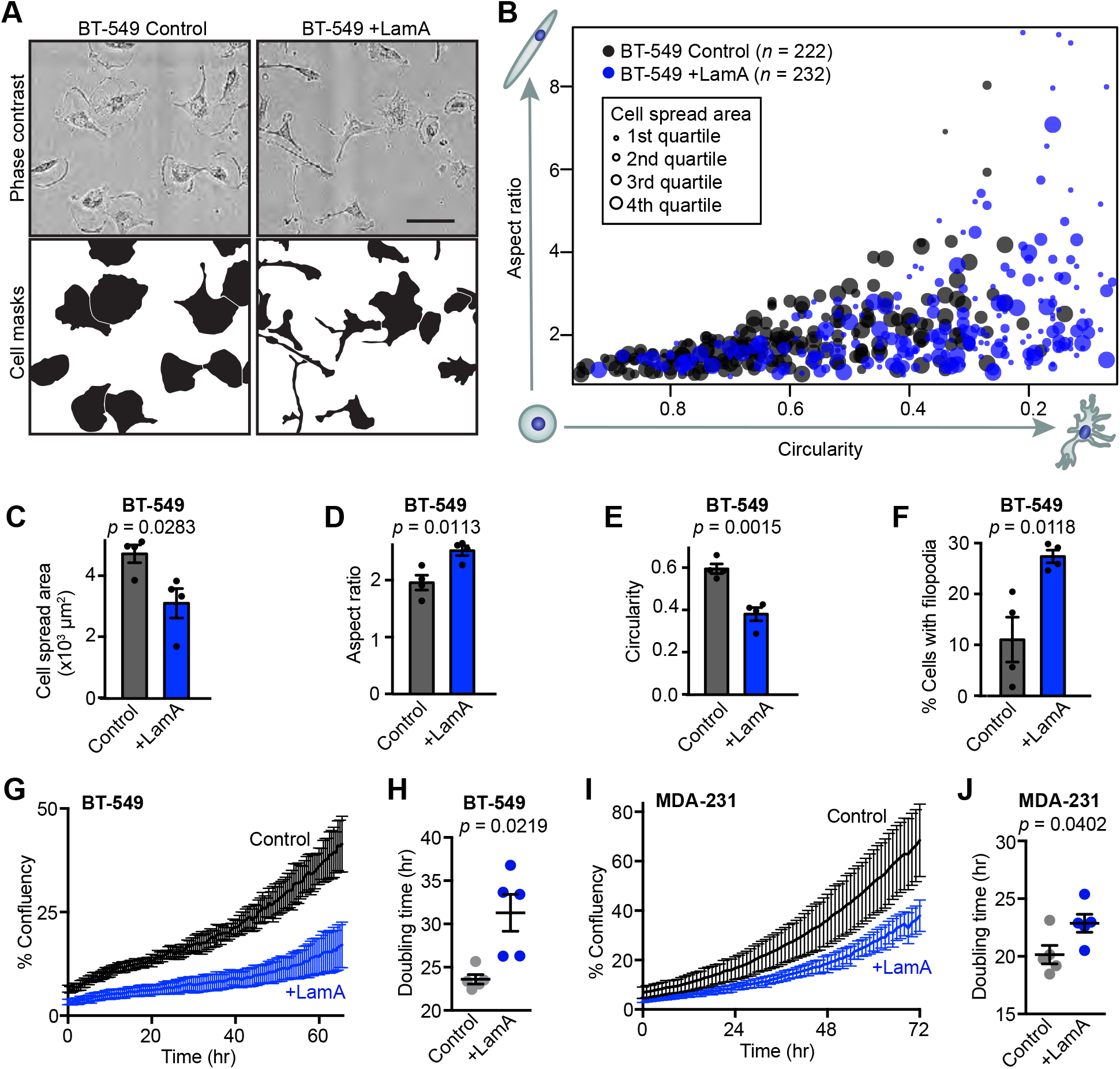
Lamin A levels modulate breast cancer cell morphology and proliferation. **(A)** Representative images showing morphology of BT-549 control and +LamA cells. Scale bar = 100 μm. **(B)** Quantification of cell morphology (circularity, aspect ratio and cell spread area) for BT-549 control and +LamA cells. Larger marker size indicates larger cell spread area. Data shown were collected across 4 independent experiments. **(C-F)** Adherent BT-549 control and +Lam A cells were quantified (*N* = 4, mean ± SEM) to determine cell spread area (C), cell aspect ratio (D), cell circularity (E), and the percentage of cells exhibiting filopodia-like protrusions (F). Statistical analysis by two-tailed unpaired Student’s *t* test. **(G)** Representative proliferation curves for BT-549 control and +LamA cells. Data shown represent a single experimental replicate where measurements were collected from images taken every 0.5 hour from 7 wells per condition plotted as mean ± SD. **(H)** Doubling times were calculated for BT-549 control and +LamA cells from *N* = 5 independent experiments. Data plotted as mean ± SEM and statistical analysis by two-tailed unpaired Student’s *t* test with Welch’s correction for unequal variances **(I)** Representative proliferation curves for MDA-231 control and +LamA cells. Data shown represent a single experimental replicate where measurements were collected from images taken every 0.5 hour from 7 wells per condition plotted as mean ± SD. **(J)** Doubling times were calculated for MDA-231 control and +LamA cells from *N* = 5 independent experiments. Data plotted as mean ± SEM and statistical analysis by two-tailed unpaired Student’s *t* test.

### Decreased lamin A/C expression is associated with increased breast cancer cell proliferation in vitro and in vivo

Since altered lamin A expression affected numerous proteins with various cellular functions, we examined whether cell proliferation was also impacted. Even a moderate (≈1.5- to 2.5-fold) increase in lamin A expression in BT-549 and MDA-231 cells, which normally have low levels of lamin A/C (Fig. 2 D, E, G, H), significantly slowed cell proliferation and increased population doubling times compared to mock controls (Figs. 6 G-J). In contrast, shRNA-mediated depletion of lamin A/C in PyMT cells, which normally express high levels of lamin A/C, did not significantly alter proliferation (Figs. S3 A and B). One interpretation of these data is that decreasing A-type lamin levels alone may not be sufficient to accelerate cell proliferation in breast cancers with initially high lamin A/C levels, but that the low lamin A/C levels found in highly aggressive breast cancers do contribute to allowing high rates of proliferation. However, we do acknowledge that many factors can contribute to altered proliferation rates in experimentally manipulated cells in culture. To test whether the link between lamin A/C levels and cell proliferation extends to human breast tumors, we assessed 21 human primary tumor tissue sections for their levels of A-type and B-type lamins in tumor cells, and for the fraction of Ki67-positive tumor cells, a marker for actively cycling cells (Fig. 7 A). Lamin A/C levels were normalized to lamin B levels to account for differences in sample processing and immunolabeling efficiency. Comparing tumors from different patients, we identified a significant correlation between the average lamin A/C levels, normalized to lamin B, and the fraction of Ki67-positive cells within the tumor (Fig. 7 B). Notably, the ratio of lamin A/C to lamin B showed a stronger correlation with the fraction of Ki67-positive cells than either lamin A/C or lamin B staining alone (Fig. 7 B and Supp. Figs. S4 A and B), highlighting the merit of normalizing the lamin A/C measurements to the lamin B signal as a control for sample processing, though this could also capture potentially meaningful shifts in relative composition of nuclear envelope components, as has been observed to occur across soft versus stiff tissues [23, 84]. Importantly, the correlation between low lamin A/C levels and high fraction of Ki67-positive cells also held true at the single cell level within individual tumors. When cells from individual tumors were split into subgroups based on their Ki67 status, the Ki67-positive subgroup had significantly lower average lamin A/C to lamin B ratios than the Ki67-negative subgroup from the same tumor (Fig. 7 C). Nearly every tumor analyzed exhibited this trend (13 of 14 tumors), demonstrating that even within a single tumor, decreased lamin A/C consistently associates with a more proliferative subset of cells. The association of lamin A with the Ki67 proliferative marker could potentially be affected by cell cycle-specific fluctuations in lamin levels, meaning that increased proliferation could be a cause of the decrease in lamin A detected rather than a consequence. This is a particularly relevant consideration since progression through the cell cycle is accompanied by phosphorylation-driven disassembly and reassembly of the lamina [85, 86]. Although we excluded mitotic cells from our analyses, we cannot rule out that cell cycle progression could impact our measurements.

**Figure 7.**
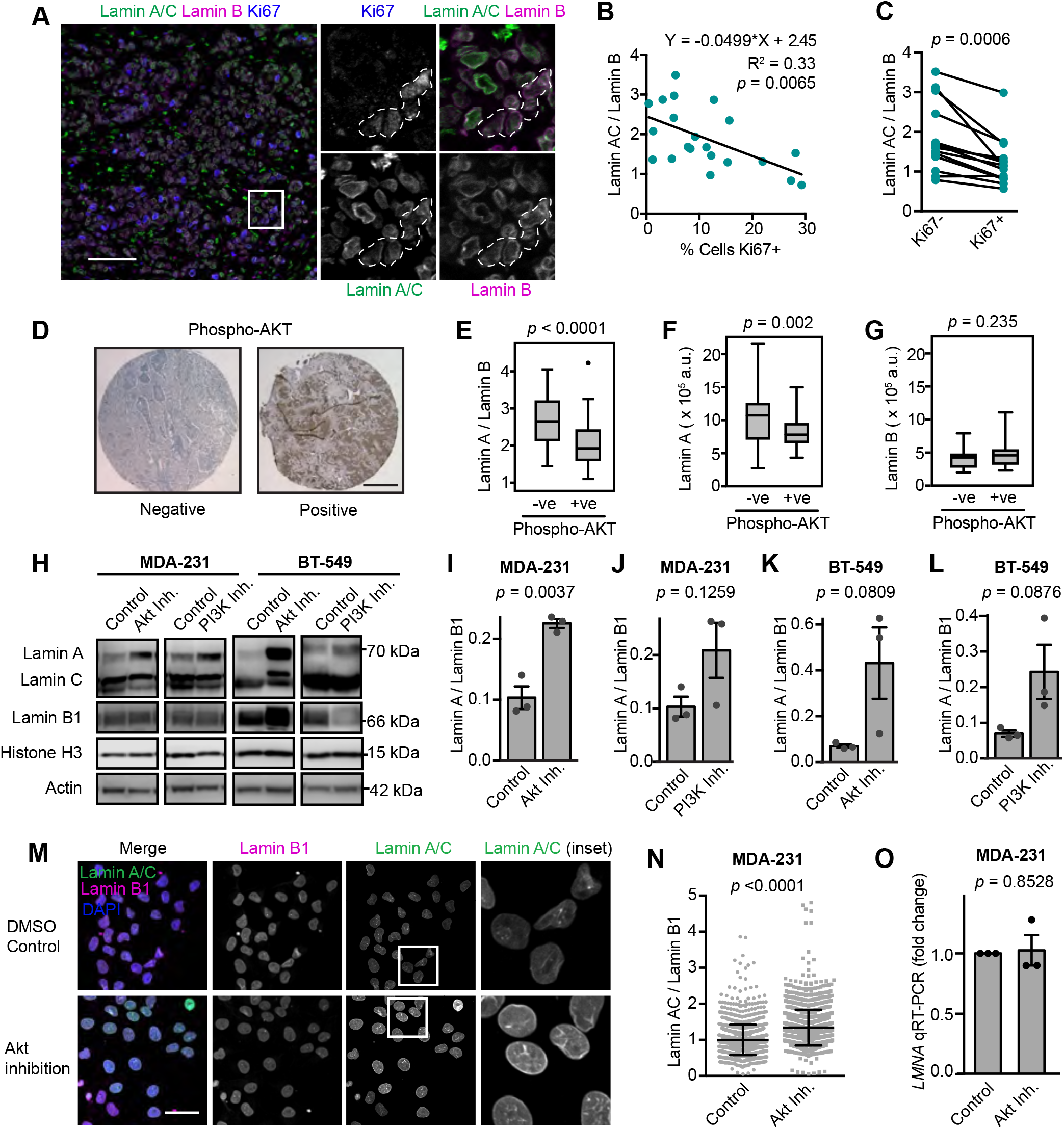
Decreased A-type lamin levels are associated with increased proliferation and Akt signaling in human breast tumors. **(A)** Images show examples of immunofluorescence staining for lamins and the Ki67 proliferative marker in a human breast tumor. Ki67-positive cells are outlined in the inset images. Scale bar = 100 μm. **(B)** The average lamin A/C to lamin B ratio per tumor (*N* = 21 human breast tumors, minimum of 76 cancer cells quantified per tumor) was quantified as nuclear rim lamin intensity from immunofluorescence staining and showed an inverse correlation with the percentage of proliferative cells. Total cell count was based on the number of DAPI-stained nuclei and cells were counted as proliferative when Ki67 positivity was 2-fold above background. The linear regression results are indicated. **(C)** Pairwise comparison of Lamin A/C to lamin B ratio in Ki67+ (proliferating) cells and Ki67– cancer cells within individual tumors (*N* = 14 tumors). Only tumors with at least 10 Ki67-positive cells were included. Lamin A/C : lamin B ratio calculated from nuclear rim immunofluorescence staining intensity. Statistical analysis by two-tailed paired Student’s *t* test. **(D)** Representative images of negative and positive immunohistochemistry staining of phospho-Akt in tumor sections from a breast cancer tissue microarray (TMA) generated from patients that had not received any systemic treatment. Scale bar = 500 μm. **(E-G)** Phosphorylation of Akt was previously scored in a breast cancer TMA [53], and immunofluorescence lamin staining and quantification of nuclear rim staining intensity in this same TMA revealed that Akt signaling was associated with lower lamin A and lower lamin A : lamin B ratio. Lamin B alone did not show a significant association with phosphorylation of Akt. Statistical analysis by two-tailed unpaired Student’s *t* test. (*N* = 89 tumors, see Wennemers et al. 2013 for original P-Akt scoring) [53]. **(H-L)** Representative Western blots and quantification of lamin levels in MDA-231 and BT-549 cells treated with Akt inhibitor (Afuresertib, 5 μM) or PI3K inhibitor (NVP-BKM120, 2 μM) for 48 hours. Histone H3 is used as a loading control (*N* = 3, mean ± SEM). Statistical analysis by two-tailed unpaired Student’s *t* test. **(M)** MDA-231 cells were treated with an Akt inhibitor (Afuresertib, 5 μM for 24 hours) and stained for lamins by immunofluorescence and representative images are shown. Scale bar = 20 μm. **(N)** Quantification of lamin immunofluorescence staining intensity in the nuclear rim in MDA-231 cells treated as in (M). Analyzed cells (*n* = 1409 and 1376 cells) were pooled from two independent experiments and plotted as mean ± SD. Statistical analysis by two-tailed Mann-Whitney test. **(O)** Analysis of *LMNA* gene expression by qRT-PCR in MDA-231 cells treated with an Akt inhibitor (Afuresertib, 5 μM for 24 hours). Data collected from *N* = 3 independent experiments and plotted as mean ± SEM. Statistical analysis by one-sample *t* test with a theoretical value of 1.

### Akt signaling promotes decreased lamin A/C

To explore which molecular mechanism may be responsible for the altered lamin A levels observed in some breast cancers, we investigated potential regulators of lamin A expression and turnover. The PI3K/Akt pathway has previously been found to influence lamin A/C expression and turnover [77, 87–89], and PI3K/Akt signaling is frequently dysregulated in cancers, where it serves as a key driver of growth and division [90]. Furthermore, analysis of SILAC proteomics data from BT-549 cells with lamin A overexpression identified the PI3K/Akt pathway in both KEGG pathway enrichment and in analysis of potential upstream regulators (Fig. 5 C and D). To explore whether the connection between Akt signaling and lamin A/C contributes to the variability in lamin A levels and nuclear deformability we observed across breast cancers (Fig. 1), we quantified lamin levels in a human breast tumor microarray (*n* = 109) that had previously been analyzed to separate tumors into groups negative or positive for phosphorylated Akt, which marks active Akt [53] (Fig. 7 D). Remarkably, tumors positive for Akt phosphorylation had significantly lower lamin A to lamin B ratios (Fig. 7 E). This was primarily driven by lamin A (*p* = 0.002, Fig. 7 F), whereas lamin B alone did not significantly correlate with p-Akt (*p* = 0.235, Fig. 7 G). To test whether Akt signaling contributes to downregulation of lamin A/C, we quantified lamin levels in BT-549 and MDA-231 cells after treatment with an Akt inhibitor (Afuresertib, [91, 92]) or a PI3K inhibitor (BKM120, [93]) (Supp. Fig. S4 C-H). Notably, both methods of targeting the PI3K/Akt pathway resulted in at least a 2-fold increase in lamin A relative to lamin B1 (Fig. 7 H-L). We obtained similar results when analyzing lamin A/C expression in MDA-231 cells treated with Akt inhibitors by immunofluorescence (Figs. 7 F and G). This increase in A-type lamin protein levels occurred without a significant increase in *LMNA* gene expression (Fig. 7 H), suggesting that Akt signaling influences lamin levels in breast cancer cells through post-transcriptional mechanisms, such as regulation of protein stability, rather than via transcriptional regulation [77, 94, 95].

### Decreased lamin A levels are associated with worse prognosis in breast cancer

Our findings suggest that primary breast cancer tumors with lower lamin A/C levels have enhanced proliferation, invasion, and confined migration abilities, which could promote cancer progression and metastasis. To investigate the proposed associations between lamin A/C levels and patient outcomes, we measured lamins levels in primary tumor tissue microarrays representing a retrospective cohort of 122 breast cancer patients, including 54 with triple-negative breast cancers, who did not receive systemic treatment and had been followed-up for at least 5 years [52]. Lamin levels were assessed using immunofluorescence and segmentation of nuclei for single cell analysis of lamin intensity at the nuclear rim, allowing measurements to reflect differences in the nuclear lamina composition regardless of nuclear morphology (Fig. 8 A). Another advantage of this approach is that nuclei can be selected so that only tumor cells are analyzed and surrounding stromal cells are excluded. The ratio of lamin A to lamin B was calculated for each individual tumor cell, and these measurements were then used to calculate the average ratio for each tumor (Fig. 8 B). Kaplan-Meier survival analysis with log-rank testing found a significant association between disease-free survival and lamin A : B ratio. Importantly, patients with breast tumors with low lamin A: lamin B ratios (based on an optimal cutoff ratio of 1.983) had significantly reduced disease-free survival compared to patients with high lamin A : B ratios (Fig. 8 C). This correlation with patient survival occurred without any significant association with ER/PR/HER2 status (Fig. 8 D), suggesting that lamin A downregulation may identify more aggressive breast cancers in a subtype-independent manner. These findings support that the impact of decreased lamin A levels on cell invasion and proliferation are functionally important for breast cancer progression, as lower lamin A levels are associated with worse disease-free survival across human breast cancer patients.

**Figure 8.**
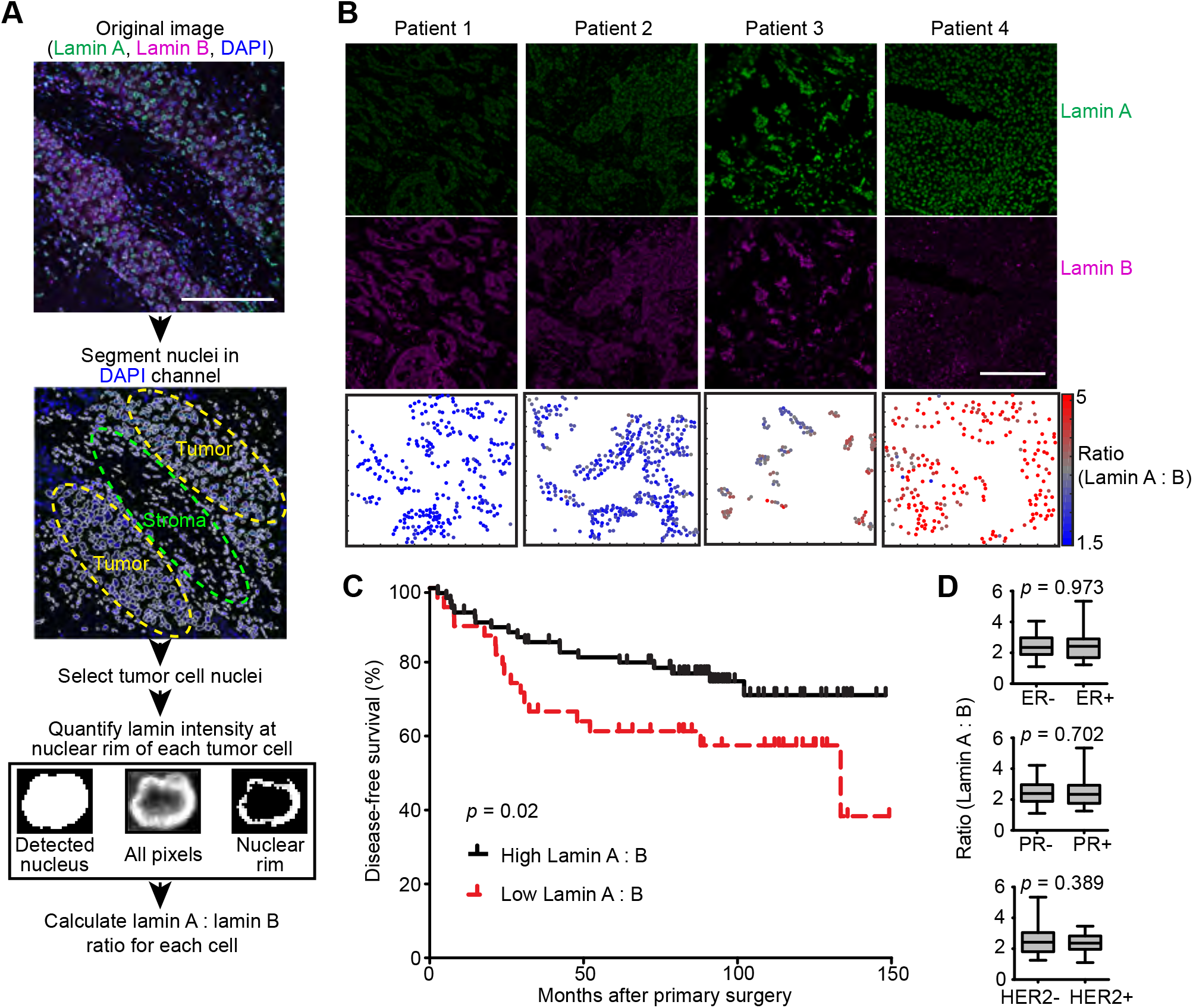
Decreased lamin A levels are associated with worse prognosis in breast cancer. **(A)** Flow chart for analysis of lamin immunofluorescence staining in human tumors. DAPI-stained nuclei were used for segmentation of individual nuclei followed by selection of regions containing tumor cells to exclude stromal cells from the analysis. Within each detected nucleus, the nuclear rim was segmented based on the top 70% of pixels above background levels. Both lamin A and lamin B intensity at the nuclear rim were quantified with this method, allowing the lamin B signal to be used for normalization of lamin A across different tumor sections. Scale bar = 200 μm. **(B)** Examples of immunofluorescence lamin staining and quantification in patient tumors. Scale bar = 200 μm. **(C)** The average lamin A : lamin B ratio per tumor was quantified from immunofluorescence staining of the TMA as shown in (A) and (B). The TMA was comprised of node-negative breast cancer patients who had not received systemic therapy and had a minimum of 5 years of follow-up. Decreased lamin A : lamin B ratio was associated with worse breast cancer patient survival as indicated by a Kaplan-Meier plot using an optimal lamin A : lamin B cutoff of 1.983 to define high (*N* = 74) and low (*N* = 35) groupings. A hazard ratio of 0.45 (95% confidence interval: 0.22-0.9, *p* = 0.02) was calculated for high lamin A: vs low lamin A: B ratio. Patient survival means were 91.6 months ± 9.1 SEM for low lamin A : B tumors and 120.9 months ± 5.9 SEM for high lamin A : B tumors. **(D)** Lamin A : B ratio was not significantly associated with estrogen receptor (ER), progesterone receptor (PR), or HER2, suggesting that lower lamin A levels identify poor prognosis independent of breast cancer histological subtype. Statistical analysis by two-tailed Student’s *t* test.

## Discussion

Despite the widespread and historic use of altered nuclear size and NE morphology for cancer diagnosis and prognosis, an integrated understanding of how these changes arise and how they relate to biochemical and mechanical functions influencing tumor progression has remained elusive. Using a broad panel of human and mouse breast cancer cells lines, we present evidence that the deformability of the nucleus varies dramatically across different breast tumor types and is determined by alterations in the levels of the nuclear envelope protein lamin A and, to a lesser extent, lamin C. Lamin B was also assessed and did not correlate with nuclear deformability, though normalizing lamin A to lamin B levels strengthened the observed relationships, indicating that the ratio of lamins A : B may provide additional insights even into processes that seem to be largely driven by modulation of lamin A. Importantly, our analysis of cell lines from isogenic and syngeneic mouse models of breast cancer found that a decrease in lamin A levels, but not lamin B, was associated with the acquisition of metastatic capabilities. Consistent with these findings, decreased lamin A levels in human tumors correlated with increased proliferation and reduced disease-free survival in breast cancer patients. Notably, proteomic and morphological analysis revealed that the effect of altered lamin A/C levels in invasive breast cancer cells extends well beyond regulation of nuclear deformability, affecting extracellular remodeling, cell adhesion, metabolism, and the PI3K/Akt signaling pathway. Our comparison of hits identified in the SILAC proteomic analysis with transcriptomic data from human tumor samples provided substantial agreement between the gene correlation with altered lamin A/C expression, suggesting that these findings are broadly applicable to breast cancer. In addition, we provide characterization of lamin A-driven phenotypes across breast cancers, and demonstrate that the oncogenic PI3K/Akt pathway is an upstream modulator of lamin A in the context of breast cancer. Collectively, these findings support that modulation of lamin A/C may have a central role in mediating broad reprogramming of cellular functions during cancer progression.

Lamin A/C expression has been previously reported to exhibit stiffness-dependent scaling as cells sense and respond to their physical environment; lamin A/C levels increase in stiffer environments and in conjunction with force transmission to the nucleus [23, 84, 96, 97]. However, we observed variations in lamin A/C in breast cancer cells that could not be explained by differences in substrate stiffness as cells were grown on identical substrates, and lamin A/C levels varied between and within patient tumors. Instead, we identified a consistent association between decreased lamin A/C and enhanced invasive, metastatic, and proliferative characteristics in cell lines and tumors. Although it is likely that multiple pathways converge to explain the variable lamin A/C expression observed in different cancers, our findings support a direct connection between oncogenic activation of PI3K/Akt signaling and the regulation of A-type lamins and nuclear deformability. Akt has previously been reported to regulate lamin A/C through both alterations in gene expression and phosphorylation-mediated protein targeting for degradation [77, 88]. The PI3K/Akt pathway is hyperactivated in ≈60% of breast cancers [98], and we found that Akt activation was associated with significantly lower lamin A : B ratios in human tumors and that Akt signaling contributes to maintaining low lamin A/C levels in aggressive breast cancer cell lines. Furthermore, we found that PI3K/Akt signaling was a significantly enriched pathway and potential inhibited upstream regulator of proteomics changes driven by lamin A expression, suggesting overlap between the pathways promoted by a decreased lamin A and the effects of Akt signaling. Thus, decreased lamin A in breast cancer may be a downstream consequence of activation of a proliferative PI3K/Akt signaling network, raising the question of whether downregulation of lamin A contributes to Akt-driven tumor progression. In support of this hypothesis, we found that a moderate increase in lamin A through exogenous expression was sufficient to decrease proliferation in aggressive breast cancer cells, and highly proliferative tumors and actively cycling cells exhibited lower lamin A : B ratios, though it is possible that cell cycle-driven fluctuations in lamin levels and/or localization contribute to these results. Our findings are consistent with previous studies linking a decrease in lamin A/C to enhanced proliferation across multiple cell types [99, 100]. Notably, multiple studies have linked lamin A/C to cell cycle regulation through interaction with pRB, thereby maintaining the ability of this tumor suppressor protein to delay cell cycle entry [101–103]. Taken together, our findings support that the downregulation of lamin A/C can be both a downstream consequence and a contributor in the reprogramming of cancer cells for enhanced proliferation.

Changes in lamina composition resulting in altered mechanical and signaling properties may also be regulated in cancers at the level of relative expression of lamin A versus C, which varies across tissues due to alternative splicing [104]. However, the majority of studies linking lamin A/C alterations to tumor progression have not distinguished between lamin A and C [39-41, 69, 105-108]. Lamins A and C are identical for the first 566 amino acids and cannot be differentially detected by most available antibodies. Interestingly, we found that the association with nuclear deformability was much stronger for lamin A than for lamin C. Furthermore, the gain of metastatic capacity in murine mammary cancer models was associated with downregulation of lamin A to a much greater extent than lamin C. These findings support that decreased levels of lamin A are the more important determinant of enhanced nuclear deformability and cancer disease progression. Consistent with this finding, decreased lamin A, but not lamin C, has been associated with metastasis and decreased disease-free survival in ovarian cancer [109], and the ratio of lamin A mRNA to lamin C mRNA has been found to decrease in multiple tumor types relative to normal tissues or less proliferative tumors [104, 110]. Accordingly, we found that aggressive breast cancer cell lines with highly deformable nuclei tend to not only have lower lamin A levels overall, but also decreased lamin A to lamin C ratios. The reasons for these isoform specific differences remain unclear. They may be related to the increased mobility and more nucleoplasmic localization of lamin C, the independent networks formed by the isoforms, or protein-protein interactions specific to their unique C-termini [111]. The potential differential roles for lamin A and C in tumor progression contrast with apparent redundant roles in tissue maintenance; mice producing only lamin C appear healthy with only minimal nuclear morphological abnormalities and a mild increase in nuclear deformability [24, 112], while mice lacking both lamin A and C die by 7 weeks of age [113]. Continued studies comparing and manipulating levels of lamin A and C isoforms are needed to better understand specific functions of individual lamin isoforms, and how changes in relative expression contribute to differentiation and disease processes.

Lamin A/C bind and regulate a multitude of proteins involved in growth factor signaling cascades, regulation of cell cycle progression, and DNA damage repair [16, 34, 114–118]. Here, we find that upon overexpression of lamin A, proteins associated with proliferative pathways and predicted inhibition of potential upstream regulators involved in proliferation (i.e., metabolism, SREBP-regulated lipid synthesis, and PI3K/Akt signaling) [119] were downregulated. One exception is the predicted activation of the EGFR family as upstream regulators, though it is important to note that while this pathway is associated with promoting proliferation early in tumor progression, it has paradoxically been found downregulated and associated with inhibition of cancer growth in metastatic cancers [120]. Lamin A overexpression also increased levels of the p53 tumor suppressor protein [121], but direct effects of p53 in this context are difficult to predict as the majority of breast cancer cell lines contain p53 mutations [122]. Nevertheless, the functional changes observed with increasing levels of lamin A do reflect characteristics typically associated with tumor suppressor proteins. However, unlike classic tumor suppressors, the *LMNA* gene is not frequently deleted or mutated in cancer [16], suggesting that a minimal level of expression is required for cell survival and growth. The balance between pathways promoted and inhibited as a function of lamin A levels may be particularly relevant to metastatic cells invading through tissues. We found that the decreased lamin A levels in highly aggressive breast cancer cells increased nuclear deformability and modulated expression of numerous proteins involved in cell adhesion and matrix remodeling, functions which are critical determinants of efficient cell movement in 3-D environments [12, 45, 123]. As a result, lower lamin A levels promote faster transit through small constrictions requiring nuclear deformation. However, squeezing the nucleus through small confinements can result in DNA damage caused by both the deformation of the nucleus and rupture of nuclear envelope [31, 124, 125]. Importantly, mutation or depletion of lamin A/C increases the frequency of nuclear envelope rupture and impairs the ability to efficiently repair and survive rupture events [27, 30, 31, 125]. The importance of maintaining some level of lamin A/C for stress resistance and epithelial cell function is further highlighted by studies showing impaired survival and growth of cells harboring severe depletion of lamin A/C [11, 31, 32, 126]. Thus, our data supports that lamin A occupies opposing roles in restricting growth and metastatic dissemination of cancer cells while also being critically important for nuclear stability, maintenance, and cell survival, as has been proposed previously [11]. Accordingly, none of the breast cancer cell lines completely lacked lamin A/C, further supporting the idea that some minimal lamin A/C expression is needed to balance promoting metastatic progression while avoiding negative effects of loss of lamin A/C.

Taken together, our data support a model where lamin A acts as a determinant of how cells interact with their physical environment, as well as a gatekeeper protecting the integrity of the nucleus in response to those interactions. Our findings support the hypothesis that primary tumors with lower lamin levels may have enhanced ability to grow and metastasize. However, this could also indicate that the low lamin A levels and highly deformable nuclei occurring in highly aggressive breast cancers are accompanied by increased sensitivity to targeted disruption of repair mechanisms involved in responses to mechanically induced nuclear damage, thus representing a potential target for novel clinical treatment options.

## Acknowledgements

We thank Prof. Peter Friedl for the 4T1 progression series cell lines. This work was performed in part at the Cornell NanoScale Science & Technology Facility, a member of the National Nanotechnology Coordinated Infrastructure, which is supported by the National Science Foundation (Grant NNCI-2025233). This work was supported by funding from the National Institutes of Health (R01 HL082792, R01 GM137605, U54 CA210184, and U54 CA193461 to J.L., R35 GM141159 and R01 GM123018 to M.B.S.), the Department of Defense Breast Cancer Research Program (Breakthrough Award BC150580 to J.L), and the National Science Foundation (CAREER Award CBET-1254846 to J.L.). The content of this manuscript is solely the responsibility of the authors and does not necessarily represent the official views of the National Institutes of Health. The results published here are in part based upon data generated by the TCGA Research Network: https://www.cancer.gov/tcga.

## Author contributions

E.S.B. and J.L. conceptualized and designed the experiments; J.L. supervised the research; E.S.B., P.S., N.Z.S., A.A.V, J.L.P.M., and A.L.M. performed experiments and analyzed data; D.K. and M.S performed the SILAC proteomic analysis; P.I., J.J.E., P.M.D, J.N.L., and V.M.W. contributed to the development of resources, including constructs, cell lines, assays, and/or image analysis methods; P.N.S. and L.V. contributed human breast tumor tissue samples and analysis; E.B. and J.L. wrote the paper; all authors contributed to the editing of the manuscript; and J.L. and M.B.S. acquired funding.

