## Supplementary Text + Figures for "Low lamin A levels enhance confined cell migration and metastatic capacity in breast cancer"

### Supplemental Figures and Figure Legends

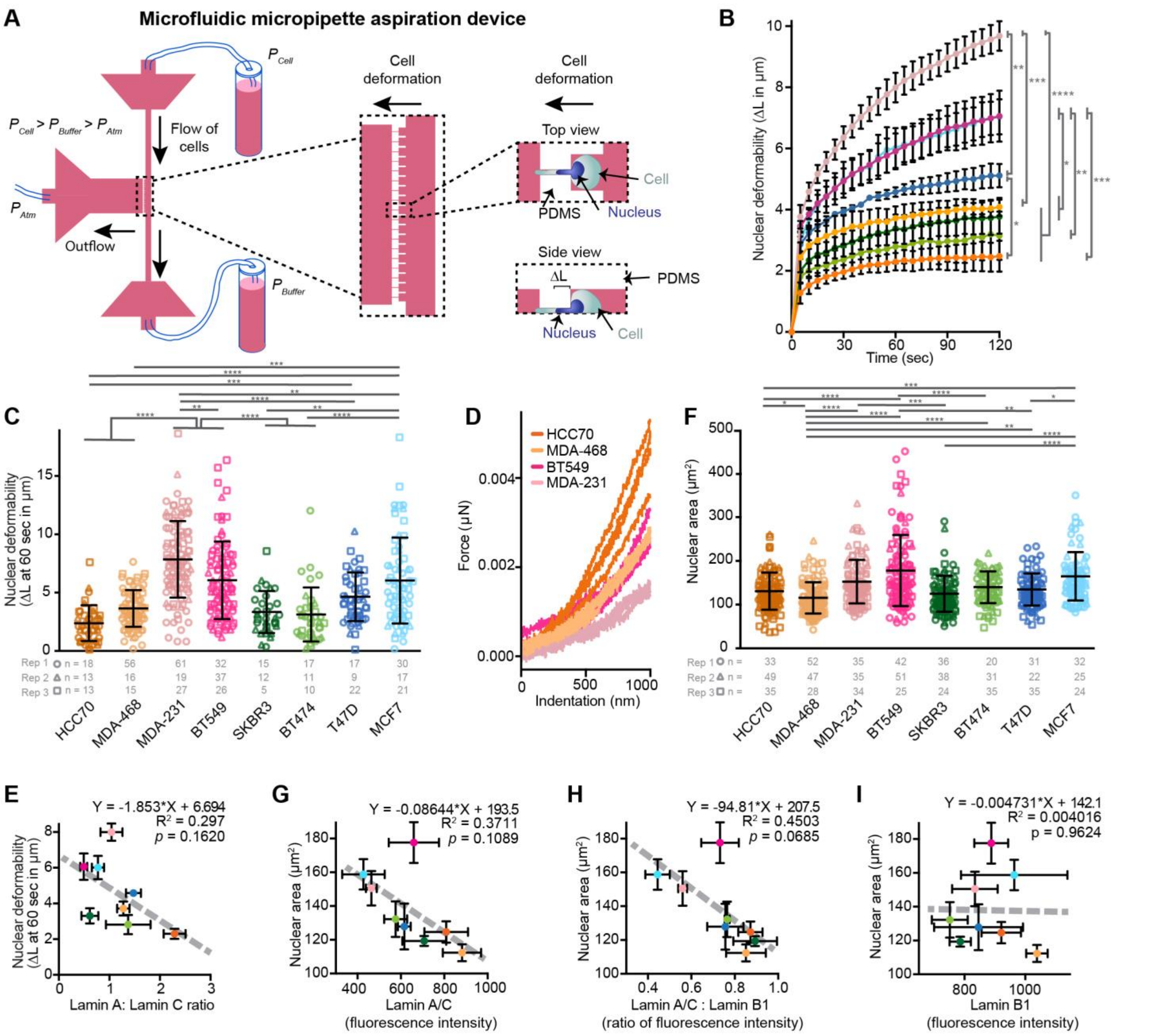


Figure S1. **Characterization of lamin levels, nuclear deformability, and nuclear area in a panel of breast cancer cell lines.** Related to Figure 1. **(A)** Schematic of the microfluidic micropipette aspiration device for high throughput assessment of nuclear deformability. A constant pressure gradient is applied across the three ports, such that cells flow into 10 × 20 μm$^2$ pockets. Nuclear protrusion (ΔL) into the 3 × 5 μm$^2$ channels is monitored over time by imaging nuclei labeled with Hoechst 33342 or another fluorescent DNA stains. Back-flushing channels manually from the outflow port allows a new set of cells to enter the channels. **(B)** Nuclear protrusion curves for eight different representative human breast cancer cell lines with different degrees of nuclear deformability. $N = 3$ independent experiments as shown in panel C, mean ± SEM, * $p < 0.05$, ** $p < 0.01$, *** $p < 0.001$, **** $p < 0.0001$. Statistical analysis by two-way repeated measures (RM) ANOVA with Tukey's multiple comparisons. MDA-231, BT549, MDA-468, and HCC70 curves are also shown in Figure 1B. **(C)** Nuclear deformability assessed based on the nuclear protrusion length into the micropipette at 60 seconds of aspiration. Individual cell measurements and number of replicates are indicated for each of 3 independent experiments. Mean ± SD, * $p < 0.05$, ** $p < 0.01$, *** $p < 0.001$, **** $p < 0.0001$. Statistical analysis by Kruskal-Wallis test with Dunn's multiple comparisons. **(D)** Representative curves from Chiaro force-indentation experiments, depicting 4 representative curves per cell type, out of a total of 34-74 cells measured per cell type, corresponding to the data in Fig. 1D. **(E)** Nuclear deformability ($N = 3$ as shown in panel C), shows an inverse correlation with the ratio of lamin A to lamin C determined by western blot analysis ($N = 3$). Data are plotted as mean ± SEM and the linear regression results are indicated on the graph. **(F)** Nuclear cross-sectional area measured on trypsinized cells in suspension in the microfluidic micropipette aspiration device. Individual cell measurements and number of replicates are indicated for each of the 3 independent experiments. Mean ± SD, * $p < 0.05$, ** $p < 0.01$, *** $p < 0.001$, **** $p < 0.0001$. Statistical analysis by Kruskal-Wallis test with Dunn's multiple comparisons. **(G-I)** Quantification of nuclear rim lamin intensity by immunofluorescence ($N = 3$ independent experiments with a minimum of 46 measurements per cell line per replicate) reveals an inverse correlation between nuclear cross-sectional area ($N = 3$ as shown in panel F) and lamin A/C levels or lamin A/C: lamin B1 ratio. Compared to lamin A/C, lamin B1 levels were less variable between cell lines and did not correlate with nuclear cross-sectional area. Data are plotted as mean ± SEM; the linear regression results are indicated on each graph.

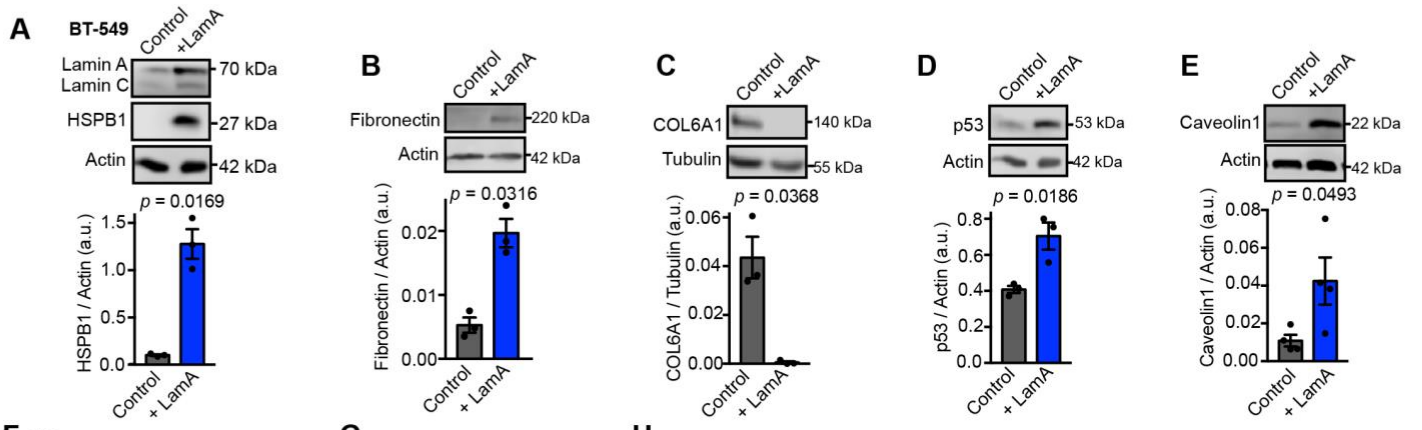


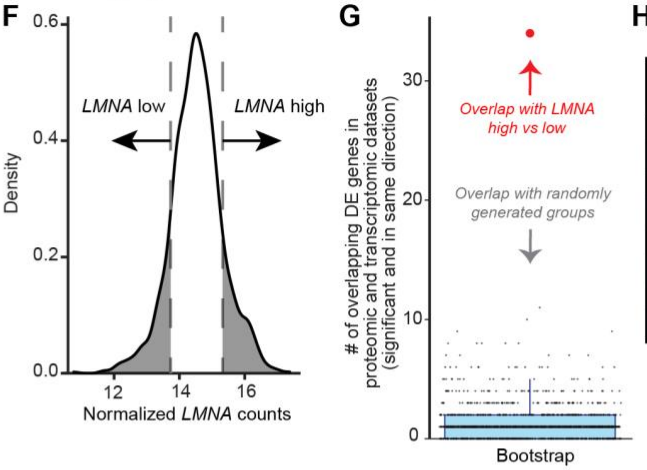


H

| KEGG Pathway | Description | Count in +LamA network | Strength | False discovery rate |
|---|---|---|---|---|
| hsa04512 | ECM-receptor interaction | 9 of 21 | 1.36 | 4.32 x $10^{-7}$ |
| hsa01100 | Metabolic pathways | 21 of 376 | 0.47 | 0.00038 |
| hsa05165 | Human papillomavirus infection | 9 of 76 | 0.8 | 0.0019 |
| hsa05165 | Protein digestion and absorption | 5 of 16 | 1.22 | 0.0023 |
| hsa04510 | Focal adhesion | 8 of 61 | 0.84 | 0.0023 |
| hsa04151 | PI3K-Akt signaling pathway | 8 of 64 | 0.82 | 0.0023 |
| hsa05205 | Proteoglycans in cancer | 7 of 58 | 0.81 | 0.0071 |
| hsa05146 | Amoebiasis | 5 of 26 | 1.01 | 0.0087 |

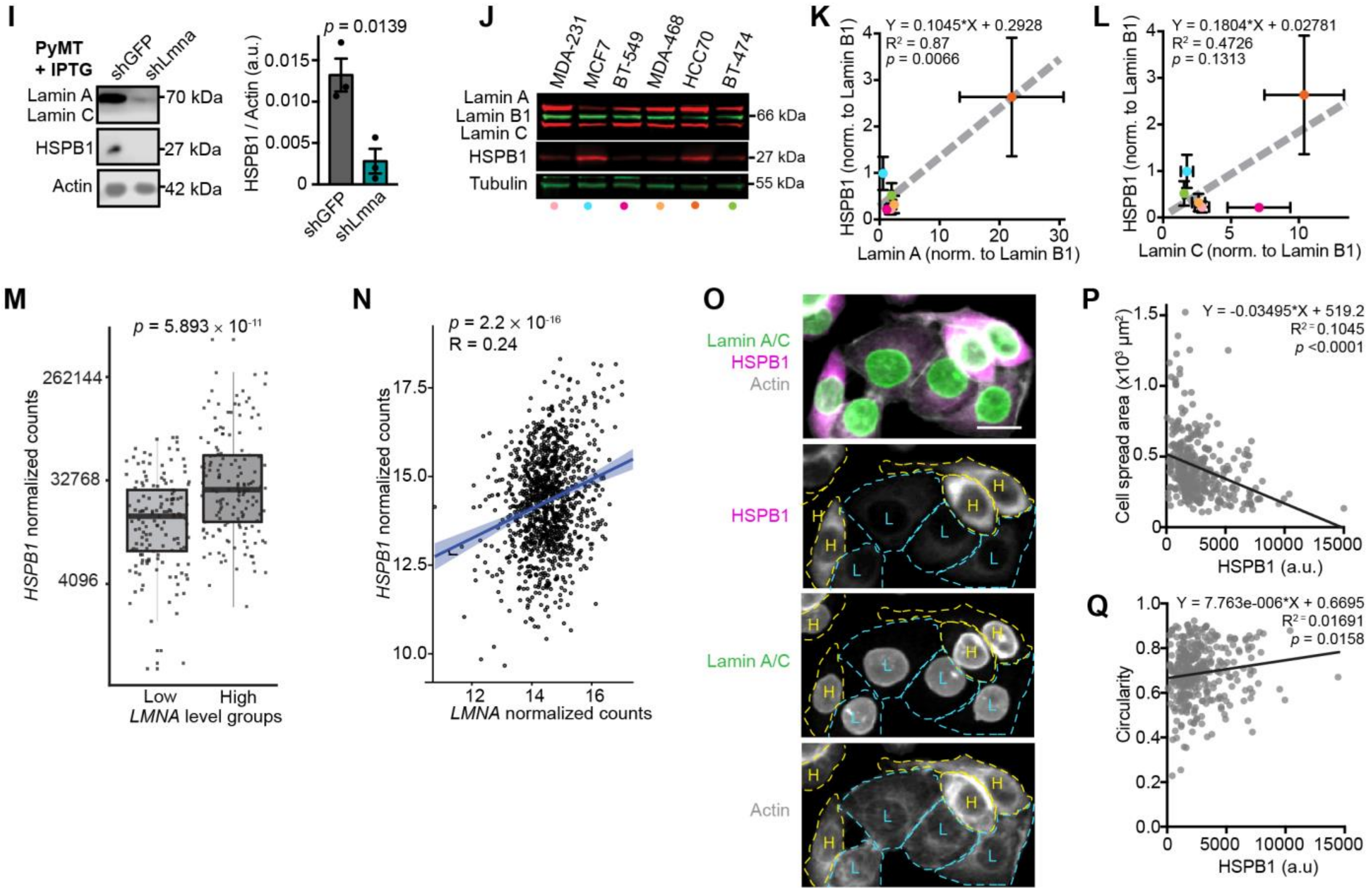

Figure S2**. Characterization of proteomic and transcriptomic changes associated with increased lamin A expression in breast cancer.** Related to Figure 5. **(A-E)** Representative western blots and corresponding quantification of protein levels in BT-549 control and +LamA cells to validate SILAC results. Relative to control cells, +LamA cells have increased levels of the molecular chaperone HSPB1 ($N = 3$). Extracellular matrix components were also altered, including increased fibronectin (FN, $N = 3$), which can be associated with wound healing and fibrosis [1], but a decrease in collagen 6A1 (COL6A1, $N = 3$), which is a protein frequently elevated in cancers and implicated in migration, Akt signaling, proliferation, and metastasis [2-6]. The tumor suppressor p53 (TP53, $N = 3$) was also increased in +LamA cells, as was caveolin 1 (CAV1, $N = 4$), a component of the plasma membrane domains involved in endocytosis, signal transduction, cell adhesion, and response to mechanical stress [7, 8]. Tubulin and actin were used as loading controls. Data displayed as mean ± SEM. Statistical analysis by two-tailed unpaired Student's *t* test. Welch's correction was applied for comparisons between groups with different variances. **(F)** Density plot of normalized RNA-seq *LMNA* counts from 1150 primary breast tumors in the TCGA database. Dashed lines indicate the edges of mean ± 1 standard deviation, highlighted regions represent "*LMNA* high" (right, $n = 153$) and "*LMNA* low" (left, $n = 145$) groups. **(G)** Differential gene expression (DEG) analysis between "*LMNA* high" and "*LMNA* low" groups showed that 34 genes (red datapoint) agreed with the direction of regulation in the SILAC proteomic comparison of +LamA BT-549 cells compared with control BT-549 cells. This was statistically significant ($p < 0.001$) as assigning the same 298 breast tumor samples to randomized groupings and running this bootstrap simulation 1000 times (black dots) returned a Poisson-like distribution with a maximum of 11 overlapping DEGs. **(H)** Full list of significant KEGG pathway enrichment for the STRING protein network generated from SILAC proteomic analysis of +LamA vs Control BT-549 cells as shown in Figure 5C. **(I)** Representative western blot and corresponding quantification ($N = 3$, mean ± SEM) of HSPB1 levels in PyMT mouse mammary tumor cells with shRNA-mediated knockdown of *Lmna* or non-targeting control shRNA. Actin was included as a loading control. Statistical analysis by two-tailed unpaired Student's *t* test. **(J-L)** Representative western blots and corresponding quantification of HSPB1 and lamin levels in breast cancer cell lines ($N = 4$). Data presented as mean ± SEM. Linear regression analysis is indicated on each graph. **(M)** Normalized counts of HSPB1 are shown for each breast tumor in the *LMNA* "high" and *LMNA* "low" groups described for panel B. Mean ± SD. **(N)** Scatter plot of normalized RNA-seq counts for *HSPB1* versus *LMNA* in 1150 primary breast tumors from the TCGA database. Blue line indicates linear regression with 95% confidence interval. Pearson correlation coefficient and *p*-values are indicated at the top of the plot. **(O-Q)** Representative images of Lamin A/C and HSPB1 immunofluorescence staining and cell shape analysis in MDA-468 cells. Cells staining strongly for HSPB1 are outlined in yellow and marked with an "H" (high) while cells with very little HSPB1 staining are outlined in cyan and marked with an "L" (low). Scale bar = 20 μm. Higher HSPB1 staining was associated with decreased spread area and more circular cell shape. Measurements of *n* = 344 cells taken from across 5 independent replicates. Linear regression analysis is indicated on each graph.

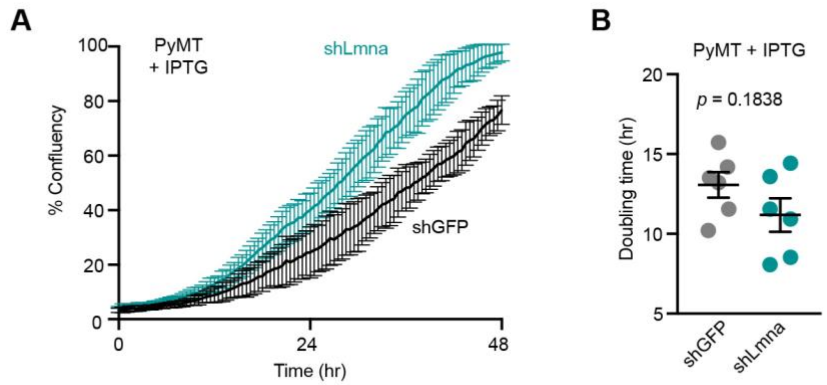


Figure S3. **Depletion of lamin A/C does not significantly alter PyMT cell proliferation.** Related to Figure 6. **(A)** Representative proliferation curves for PyMT cells with IPTG-inducible lamin A/C depletion (shLmna) or shGFP control. Data shown were acquired from a single experimental replicate where measurements were collected from images taken every 0.5 hour from 3 wells per condition and plotted as mean ± SD. **(B)** Doubling times were calculated for PyMT IPTG-inducible sh*Lmna* and shGFP control cells from $N = 6$ independent experiments. Data plotted as mean ± SEM and statistical analysis by two-tailed unpaired Student's *t* test.

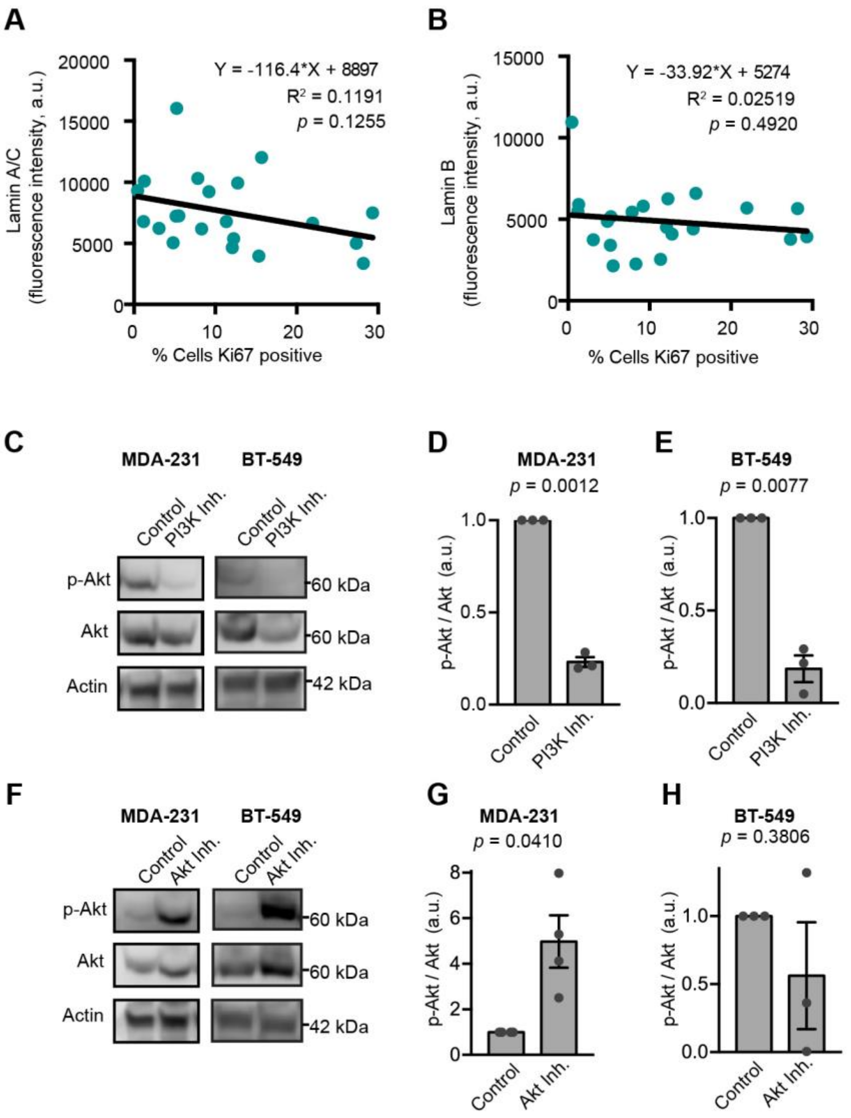


Figure S4. **Relationship between lamin levels, proliferative index, and Akt activation in human breast tumors.** Related to Figure 7. **(A-B)** The average Lamin A/C and lamin B nuclear rim staining intensity was quantified per tumor ($N$ = 21 human tumors) and assessed for correlation with the percentage of proliferative cells as determined by quantification of immunofluorescent staining of Ki67. Total cell count was based on the number of DAPI-stained nuclei and cells were counted as proliferative when Ki67 positivity was 2-fold above background. The linear regression results are indicated on each graph. **(C-E)** Representative Western blots and quantification of Akt phosphorylation in MDA-231 and BT-549 cells treated with the PI3K inhibitor NVP-BKM120 (2 μM for 48 hours). As expected, inhibition of PI3K reduced p-Akt levels. Actin is shown as a loading control and data were normalized to p-Akt/Akt in vehicle (DMSO)-treated control cells ($N$ = 3, mean ± SEM). Statistical analysis by one-sample $t$ test with a theoretical value of 1. **(F-H)** Representative Western blots and quantification of Akt phosphorylation in MDA-231 and BT-549 cells treated with the Akt inhibitor, Afuresertib (5 μM for 48 hours). Akt inhibition with Afuresertib is known to cause accumulation of hyperphosphorylated Akt but suppresses phosphorylation of Akt substrates [9, 10]. Actin is shown as a loading control and data were normalized to p-Akt/Akt in vehicle (DMSO)-treated control cells ($N$ = 4 for MDA-231 and $N$ =3 for BT-549). Data represented as mean ± SEM. Statistical analysis by one-sample $t$ test with a theoretical value of 1.

### Supplemental Video Legends

Video 1. **Nuclear deformability of MDA-231 and HCC70 cells in a micropipette aspiration microfluidic device, related to Figure 1.** Time-lapse fluorescence microscopy showing deformation of MDA-231 (left) and HCC70 (right) nuclei fluorescently labeled with Hoechst 33342 (cyan in merge, white in single channel images). Images were captured at 5 second intervals and played at 5 frames per second. Scale bar = 50 μm

Video 2. **Migration of MDA-468 shLMNA and shControl cells in a microfluidic device, related to Figure 3.** Time-lapse fluorescence microscopy showing migration of MDA-468 depleted for lamin A/C via shLMNA (right) and shControl cells (left) cells through 2 × 5 $\mu m^2$ constrictions in a microfluidic device in a 0.1% to 12% FBS gradient. MDA-468 nuclei are visible due to expression of NLS-GFP (green). Images were captured at 10-minute intervals and played at 5 frames per second. Scale bar = 50 μm

Video 3. **Migration of MDA-231 Control and lamin A overexpressing cells in a microfluidic device, related to Figure 3.** Time-lapse fluorescence microscopy showing migration of MDA-231 control cells (left) and cells with exogenous expression of lamin A (“+LamA”, right) through 2 × 5 $\mu m^2$ constrictions in a microfluidic device. Cells express GFP (green). Images were captured at 10-minute intervals and played at 6 frames per second. Scale bar = 50 μm
